# Tunable 3D Alveolosphere Model from Human Alveolar Cells: A Breakthrough Tool to Explore Emphysema Pathophysiology

**DOI:** 10.1101/2025.04.09.645486

**Authors:** Marina Guecamburu, Arthur Pavot, Caroline Jeannière, Yaniss Belaroussi, Matthieu Thumerel, Emma Sammaniego, Hugues Begueret, Guillaume Maucort, Fanny Decoeur, Jean-William Dupuy, Anne-Aurélie Raymond, Pauline Esteves, Leo Grassion, Gael Dournes, Patrick Berger, Elise Maurat, Katharina Raash, Eloïse Latouille, Vincent Studer, Isabelle Dupin, Pauline Henrot, Maéva Zysman

## Abstract

**Rationale:** Three-dimensional (3D) culture models such as alveolosphere, provide a unique tool to study emphysema mechanisms. Reducing the heterogeneity of alveolospheres which are currently mostly grown in Matrigel remains challenging.

**Objectives:** To develop a tunable and reproducible 3D-alveolosphere model exclusively from human primary type II alveolar epithelial cells (AEC2) for modeling and understanding emphysema.

**Methods:** AEC2 cells (HTII-280+) were isolated from 52 smoker and non-smoker lung samples and cultured in preformed photopolymerized hydrogel microwells of adjustable shape and size. Topological and phenotypic characterization were performed at Day (D)1, 7 and 14. Lamellar bodies (LB) were quantified using artificial intelligence (AI)-based image analysis of transmission electron microscopy (TEM) serial block face images. Emphysema was modeled through chronic exposure to 1 and 5% cigarette smoke extract (CSE) during 5 consecutive days.

**Measurements and Main Results:** 3D-alveolospheres were maintained in culture for 14 days with central lumen formation observed from D7 to D14. Presence of tight junctions (TEM imaging and ZO-1 immunostaining) suggested epithelial barrier formation. AEC1 markers (*p2xr4, pdpn*) appeared progressively from D1 to D14 while AEC2 markers (*abca3, sftpa, sftpc*) persisted over time. TEM images indicated surfactant synthesis (LB, lipid bodies) and AI-driven LB quantification showed a decrease in the proportion of LB-containing cells over time. CSE exposure led to cell death, architectural disorganization, oxidative stress and inflammation.

**Conclusion:** This standardized and adjustable 3D-alveolosphere model from human primary AEC2, reproduced key native alveolar features. CSE exposure provides an opportunity to appropriately study the pathophysiological pathways involved in emphysema.

## Introduction

Emphysema is a severe and worldwide prevalent respiratory disease due to long-term exposure to inhaled particles such as cigarette smoke [1]. It is a major feature of chronic obstructive pulmonary disease (COPD), resulting from alveoli tissue damage, causing breathlessness and altered quality of life. Anatomically, it corresponds to lung alveoli destruction, resulting in airspace enlargement and decrease in elastic recoil, due to an imbalance between protease and antiprotease activities, enhanced apoptosis, lung repair failure and/or oxidative stress [2]. Adult alveoli contain bipotent stem cells called type II alveolar epithelial cells (AEC2), which can self-renew and differentiate into type I alveolar epithelial cells (AEC1), which account for the vast majority of alveolar surface.

While animal models have paved the way for our understanding of pathobiology and the development of therapeutic strategies for COPD management, their translational capacity is limited, especially concerning emphysema modelling with cigarette smoke exposure in rodents. Therefore, there is a well-recognized need for innovative *in vitro* models built with human cells to better reflect COPD physiopathology. Three-dimensional (3D) models, especially organoids defined as auto-assembly of these cells to a 3D culture grown from pluripotent stem cells (PSCs) or adult progenitors, provide a unique tool to model alveolar functions [3, 4], such as barrier function and surfactant production [5], which are both impaired in patients with emphysema [6, 7]. Alveolospheres generated by embedding alveolar progenitors (AEC2) into a synthetic matrigel© which self-organize into spheres [5] of varying sizes and shapes, show heterogeneity in size, shape and composition that alters the robustness of these models and represents a major limitation in their use to understand disease development and progression. Reducing this physical variability [8, 9] and replicating *in vivo* alveoli anatomy, including lumen formation, remains a major challenge [10].

Microstructured hydrogels could provide a stringent template to control organoid size [11], but they are often moulded on commercial surfaces, limiting modulation of microwells shape and substrate rigidity. Therefore, tailoring the structural properties of the hydrogel for alveolospheres’ development by combining spatially tunable sizes, shapes as well as stiffness would give an unprecedented tool to increase reproducibility. In the present study, we were able to form alveolospheres by seeding human primary AEC2 (HTII-280+) into microwells of standardized size, shape and stiffness, produced by the PRIMO© technology. We demonstrated progressive lumen formation and the presence of AEC2 and 1 through innovative methodology using transmission electron microscopy and AI-driven image quantification. In addition, we provided evidence for reproducing emphysema features (such as oxidative stress, inflammation and cell junction alterations) following chronic cigarette smoke extract exposure.

## Methods

### Production of microwells

Microwells manufacturing was performed using the PRIMO system© marketed by Alveole^™^company as previously published [12]: four 3×3 mm polydimethylsiloxane (PDMS) microwells (Alvéole Stencell) were deposited on a 35mm diameter Ibidi® glass bottom Petri dish of adjustable shape, size and stiffness. Polyethylene glycol acrylate powder (4ARM-PEG-AVRL-10K, hydrogel of a semi-rigid nature) and photoinitiator solution (PLPP, benzophenone gel) were mixed in PBS and then added inside each PDMS microchamber. Finally, exposure to patterned UV irradiation (30 mj/mm) by a device comprising digital micromirrors to project the desired patterns (Alvéole PRIMO) allowed hydrogels’ photopolymerization and obtention of tailor-made micro-wells (54 microwells in each microplate) (**Figure 1A**). Stiffness of hydrogel was evaluated by microindentation (Chiaro, Optics11). Stiffness was modulated by tuning the UV dose during the photopolymerization of the hydrogels to obtain the same stiffness as described in emphysema *versus* control native lung, e.g. 1 *versus* 5kPa [13–15](**figure 1B**).

**Figure 1.**
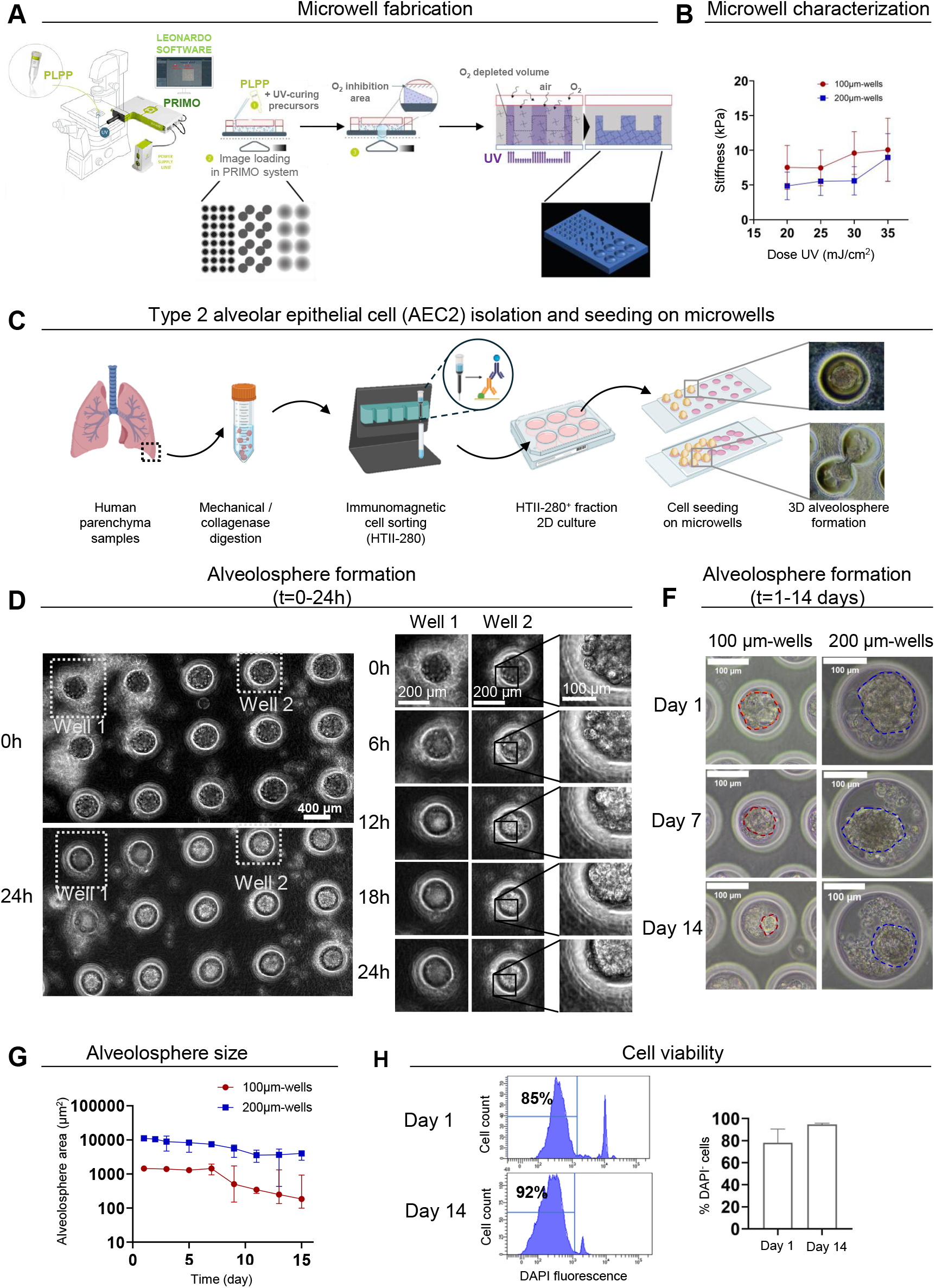
3D-alveolosphere formation from primary human type 2 alveolar epithelial cells. (A) Formation of the hydrogels microwells by UV transillumination. Light-based toolbox is able to craft microniches of adjustable shape, size and stiffness, in hydrogel thanks to photoinitiator solution (PLPP, benzophenone gel) and their interaction with oxygen. (B) Microwells’ stiffness, measured by Chiaro Optics 11, for 100 and 200µm diameter-wells. (C) Isolation and culture of type 2 alveolar epithelial cells (AEC2) obtained from human lung samples into microwells of different shapes for alveolosphere formation. AEC2 were purified using HTII-280 (Human type 2 cells-280kDa protein) immunomagnetic-activated cell sorting beads. (D) Bright-field live imaging of AEC2 seeded into microwells during the first 24h. Left panels: representative images at the initial state of the acquisition (0h) and 24 hr after cell seeding. Right panels: representative magnified images of Well 1 and Well 2 (shown in the right panel). (F) Representative bright-field images of AEC2 seeded into microwells of different sizes (diameters : 100 and 200µm) for a period of 14 days.The dotted lines indicate the contour of the alveolospheres. (G) Quantification of alveolosphere areas over time in microwells of 100 and 200µm-diameters (200µm-diameter: n=5 experiments, 100µm-diameter n=3 experiments). (H) Viability of alveolospheres grown in 200µm-diameter microwells quantified by flow cytometry Left panel: histograms showing representative cell counts (y-axis) versus DAPI fluorescence (x-axis) day 1 and day 14 after AEC2 seeding. The percentages indicate the number of DAPI-cells. Right panel: percentage of DAPI-cells day 1 and day 14 after AEC2 seeding (n= 3 experiments). 2D: 2-dimension, 3D: 3-dimension, AEC2: alveolar epithelial cell, HTII-280: Human type 2 cells-280kDa protein, t: time

*Lung tissue processing* AEC2s were isolated from tumor-free lung tissue obtained from patients undergoing surgery for lung cancer (smokers, ex-smokers, and non-smokers) or from emphysematous lung explant after transplantation at the Bordeaux University Medical Center (France) (**Suppl Figure S1**). Patient characteristics are summarized in **Table 1**. We used samples of more than 0.7 grams dissociated mechanically and enzymatically (**suppl methods**).

### AEC2s selection and culture

The AEC2s were isolated using immunomagnetic cell sorting based on the surface expression of human type 2 cells-280kDa protein (HTII-280, Terrace Biotech, **suppl methods**). AEC2 cells were initially expanded in 2D 6-well plates (seeding concentration 500,000 cells/well) in Pneumacult ExPlus culture medium (Stemcell technologies). After 5 to 7 days, the medium was changed to Pneumacult Ex basal culture medium (Stemcell Technologie) and cells were further amplified until reaching 70 to 90% confluency (**Figure 1C**).

### Seeding of AEC2 in 3D-microwells

AEC2 cells were incubated with 0.05% trypsin (Trypsin-EDTA Solution, Gibco) and reseeded in 3D seeding culture medium (Pneumacult Airway Organoid Basal Medium, StemCell) at the concentration of 10,000 cells/µL (**Figure 1C**). Forty-eight hours later, the culture medium was replaced by Organoid differentiation medium (Stem Cell). Medium was then changed every 3 days and alveolospheres were photographed daily with a brightfield microscope until day 14.

### RNA Isolation, cDNA, and quantitative Polymerase Chain Reaction (qPCR) Analysis

At each time point of interest, 350µL of RLT and βmercapto-ethanol (BME) lysis buffer (Qiagen RNeasy kit (Qiagen, Hilden, Germany)) were added to two alveolospheres’ dishes, following the manufacturer’s recommendations. The lysates were then stored at -80°C. Ten to 20 mg of lung tissue was used as a reference to normalize the pPCR results obtained at D1, D7, and D14. For the purification step, the “Qiagen RNeasy kit protocol” was used as recommended (**suppl table 1**).

### Transmission electron microscopy

3D alveolospheres were analyzed by transmission electron microscopy and serial scanning electron microscopy (Serial Block Face, SBF-TEM) as detailed in the **suppl. methods** of a whole organoid is analyzed, volumes can be reconsctructed, revealing the 3D architecture of organelles inside.

### Artificial intelligence-driven structure prediction for lamellar bodies in SBF-TEM

We also developed an artificial intelligence (AI)-driven structure prediction for lamellar bodies (LB) in whole organoid samples. Automated cell segmentation was trained by AP on diverse and previously annotated datasets by MG, MZ and AP. Briefly, following cell segmentation *via* cellpose©, morphological features such as nucleus and LB recognition has been extracted *via* Ilastik©. Hence, we applied AI-driven automated quantification to assess architectural changes over time (**suppl. methods and suppl figure 2**), especially concerning the number of cells with LB (considered to be AEC2), as compared to those without LB (considered to be AEC1).

### Flow cytometry

Lung tissue samples were dissociated with 0.05% trypsin–EDTA (Gibco™) and cell-surface antigen staining was performed by incubating cells with primary antibodies coupled to appropriate fluorophores diluted at 1:25 for 60 min at room temperature. List of antibodies can be found in (**suppl. methods and suppl table 2)**.

### Exposure to cigarette smoke extract

The cigarette smoke extract (CSE) 100% was generated as follow: the smoke from 2 research-grade filtered cigarettes (Kentucky tobacco research and development center at University of Kentucky; Lexington, KY) was bubbled over 20mL of DMEM medium (Stable Cell DMEM-high glucose, Sigma Aldrich) using a disposable tube and a manually operated syringue then sotred at -80°C. The CSE was then thawed upon use and diluted to 1 or 5% with Organoid Differenciation culture medium, accordingly. Alveolospheres were exposed since day 7, for 5 consecutive days to CSE and medium was replaced every day. Cell viability after 5 days of exposure was studied qualitatively by calcein labelling (Calcein AM, Thermofisher, C3099).

### Label-free quantitative proteomics

Cells lysis were processed using RIPA buffer. Each lysate was centrifuged and the supernatant was used for the proteomic analysis (**suppl methods)**. The mass spectrometry proteomics data have been deposited to the ProteomeXchange Consortium via the PRIDE (X3)[16] partner repository with the dataset identifier PXDxxxxx.

### Emphysema quantification on CT-scans

Clinical chest CT (GE Revolution®) of patients were collected and emphysema was analysed (**suppl methods and suppl. figure 3)**. Briefly, the proportion of lung voxels with an attenuation lower than -950 Hounsfield Units (LAA-950HU) is defined as emphysema and must be considered significant when higher than 6% [17].

### Ethics and statistics

The use of surplus lung tissue for research following surgery was within the framework of patient care, and tissue donation was based on a no-objection system for coded anonymous use of waste tissue, left-over from diagnostic or therapeutic procedures. This non-interventional research protocol was approved by the Bordeaux University Hospital’s TUBE agreement, version 1 (CHU BX 2020/54, 14/01/2021). All methods were carried out in accordance with relevant guidelines. Continuous variables are presented as means +/-standard deviation or medians [Q1-Q3] and compared using non-parametric tests (Wilcoxon, Kruskal-Wallis, Spearman). Data are depicted as individual data points with standard deviation and significance was tested using a paired t test, and differences were considered significant at p<0.05. Analyses were performed using ImageJ, ImageLab and Statistical analysis was performed using Graphpad Prism version 10 (GraphPad Software Inc. La Jolla, CA).

## Results

### Generation of standardized alveolospheres from human primary alveolar type 2 cells (AEC2)

Seventy-two lung samples were collected between 1^st^ February 2023 and 31^st^ July 2024. After the exclusion of 20 samples because of insufficient initial AEC2 amplification in 2D culture (n=14), lack of cellular adhesion to hydrogel (n=3) or contaminations (n=3), 52 lung samples were included in the analysis (**Suppl Figure S1)**. Median weight of parenchymal tissues was 2.8 [1.4; 3.9] grams and the median 2D amplification phase duration for AEC2 was 25 [20; 27] days. Patients’ characteristics are summarized in **Table 1 and Suppl Tables 3 and 4**. Briefly, 52% of them were female, with a median age of 65 [58; 71] years old, 36 (69%) were former smokers and the median smoking was 29 [10; 50] pack-years. Six (12%) patients had an airflow obstruction (defined by forced expiratory volume in 1 second (FEV1)/ forced vital capacity (FVC) < 0.7) and 4 (8%) had significant emphysema defined by a LAA-950UH above 6% measured by CT scan (**Suppl Figure S3**). Most of the samples (82%) came from lobectomy for lung cancer. There was no significant association between the number of purified AEC2 per lung sample weight and airflow obstruction nor 2D culture duration or between 2D culture duration and airflow obstruction. Furthermore, there was no correlation between the number of AEC2/g and smoking history in pack years or age. However, the failure of the amplification step in AEC2 2D culture was more frequent in patients with airflow obstruction (**Suppl Table S5**).

HTII-280 cell sorting led to an enrichment of epithelial cells into the positive fraction, while no epithelial cells were detected into the negative fraction (**Suppl Figure S4**). Cell viability remained high during cell purification, except just after cell sorting in the positive fraction (**Suppl Figure S4**). The 2D culture step enabled both the amplification of viable epithelial cells and the elimination of the contaminating CD45+ cells of the positive fraction: just before seeding into the microwells, the population was 96% DAPI negative and 84 % pan-cytokeratin positive (**Suppl Figure S4C and E**). After 20 +/-5 days in 2D culture, cells still expressed epithelial markers (pan-cytokeratin, E-cadherin) and are negative for hematopoietic markers (CD45). In addition, FACS indicated a high proportion of epithelial cell adhesion molecule (EpCAM) positive cells (**Suppl Figure S5 A**); they expressed surfactant protein C (SPC) and were positive for lysotracker, but negative for podoplanin (PDPN) and HTI-56 (human Type 1 Cells) stainings (**Suppl Figure S5 B-D**), also indicating that these cells were mainly AEC2.

Then tailored microwell allowed to test different sizes, shapes and rigidity of microwells for alveolospheres culture (**Figure 1 A, B and E**). After seeding into microwells, self-compaction of AEC2 into spherical structures was evidenced over the 24 first hours (**Figure 1 D, E**). The alveolospheres were then followed in culture for 14 days (**Figure 1 F, G**). We obtained an overall 72% success rate for the formation of alveolar organoids from successfully amplified AEC2 (**Suppl Figure S1**). The diameter of alveolospheres were higher when grown in 200 µm diameter wells compared to 100, 150 and 250 µm diameter wells (**Figure 1F, G, Suppl Figure 6 A, B**). Also, cell viability assessed by flow cytometry, 86 and 95% at day 1 and 14, respectively, confirmed the sustainability of our model (**Figure 1 H, I**). Tailor-made microwells, such as double sphere shape, were also feasible and cells grown in such microwells were viable till 14 days (**Figure 1C, Suppl Figure 6 A**). Thus, considering both human alveolar size (≈200µm) *in vivo* [18] and above-mentioned results, we chose to use 200µm diameter microwells for the rest of the experiments.

### 3D model characterization: topology, functionality

The progressive formation of the lumen, assessed by hematoxylin-eosin (HE) coloration on organoid sections, 3D confocal imaging of the whole structure and TEM imaging, was a major feature confirming the self-organizing ability of AEC2 cells into alveoli-like 3D models (alveolospheres) (**Figure 2 A-C**). By HE staining performed on sections of paraffin-embedded samples and TEM imaging, we evidenced the apparition of several internal cavities around day 7, which merged into a single central lumen around day 14, resulting in an architecture similar to that found *in vivo* **(Figure 2 A, C)**. We confirmed the presence of an internal lumen by performing 3D imaging by confocal microscopy (**Figure 2 B**). We next evaluated markers of epithelial barrier, focusing on Zona Occludens-1 (ZO-1) expression and junction characterization by electron microscopy. AEC2 became cohesive and adherent over time and we observed tight and adherens junctions assessed by both immunofluorescence (**Figure 2 D**) and TEM (**Figure 2 E**), at D7. The tight junctions and microvilli were located at the apical part of the AEC2, inside the lumen (**Figure 2 D-E**), suggesting the formation of a partially polarized epithelium.

**Figure 2:**
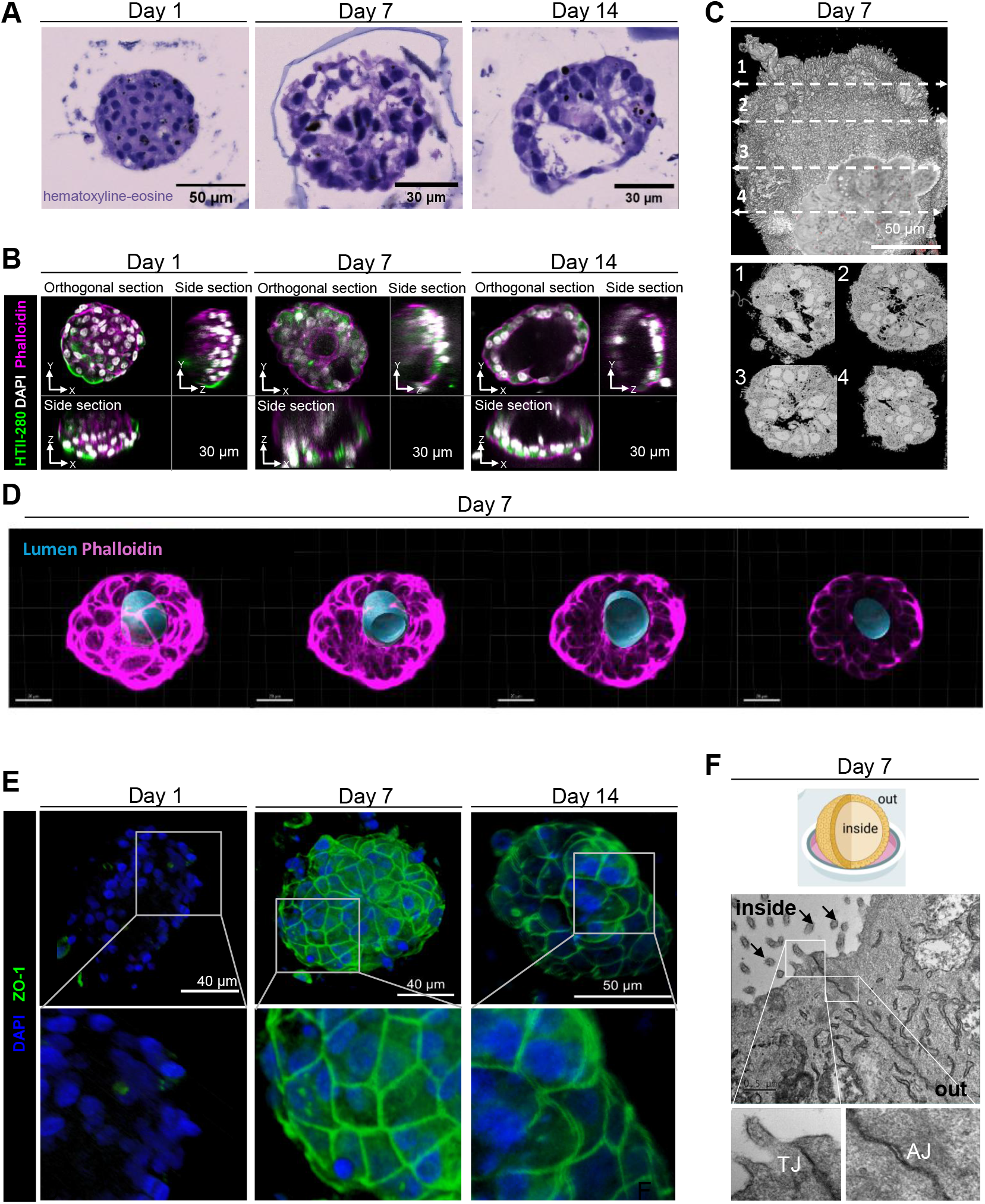
Topology of the 3D-alveolosphere model: lumen formation, barrier function, and polarity. (A) Representative bright-field images of hematoxyline-eosine (HE)-stained sections at days 1, 7, and 14. (B) Fluorescent images of alveolospheres stained for F-actin (phalloidin, magenta), the marker of type 2 alveolar epithelial cells (AEC2s) (Human type 2 cells-280kDa protein (HTII-280), green) and nuclei (DAPI, white) showing progressive lumen formation and the persistence of AEC2s at days 1, 7 and 14. (C) Reconstruction of a complete alveolosphere imaged by spinning disk Live SR microscope, at day 7 from serial sections showing lumen at the center of the organoid. (D) Transverse sections of a three-dimensional (3D) alveolosphere obtained from Z-stack spinning disk images of a 7-day-old alveolosphre stained for F-actin (phalloidin, magenta) and lumen reconstruction (blue) (E) 3D reconstruction of an alveolosphere stained for Zona Occludens-1 (ZO-1, green) and nuclei (DAPI, blue) and imaged by spinning disk microscopy at days 1, 7 and 14, showing progressive tight junction formation (TJ). (F) Top panel: schematic of an alveolosphere grown in microwell with inside-out orientation. Created with BioRender.com. Bottom panels: electron microscopic image of an embedded alveolosphere at day 7 showing intercellular adhesion between cells, with tight junctions and adherens junctions (AJ). Arrows indicate microvilli.

### Characterization of cellular differentiation

Phenotypic characterization was performed over time. The qPCR results showed the persistence of AEC2 marker gene expression (*ABCA3, SFTPA, SFTPC*) over time, while markers of intermediate cells (*CLDN4*) and AEC1 cells (*P2×4R, PDPN*) increased progressively (**Figure 3 A**). Pancytokeratin staining confirmed epithelial cells (**Figure 3 B)**. HTII-280 (**Figure 2 B**) and thyroid transcription factor-1 (TTF-1) (**Figure 3 C**) immunostainings confirmed the persistence of AEC2 cells at D1, 7 and 14. Finally, presence of surfactant-producing organelles was also evidenced through LysoTracker staining (**Figure 3 D**), which has been shown to accumulate in LB [19].

**Figure 3:**
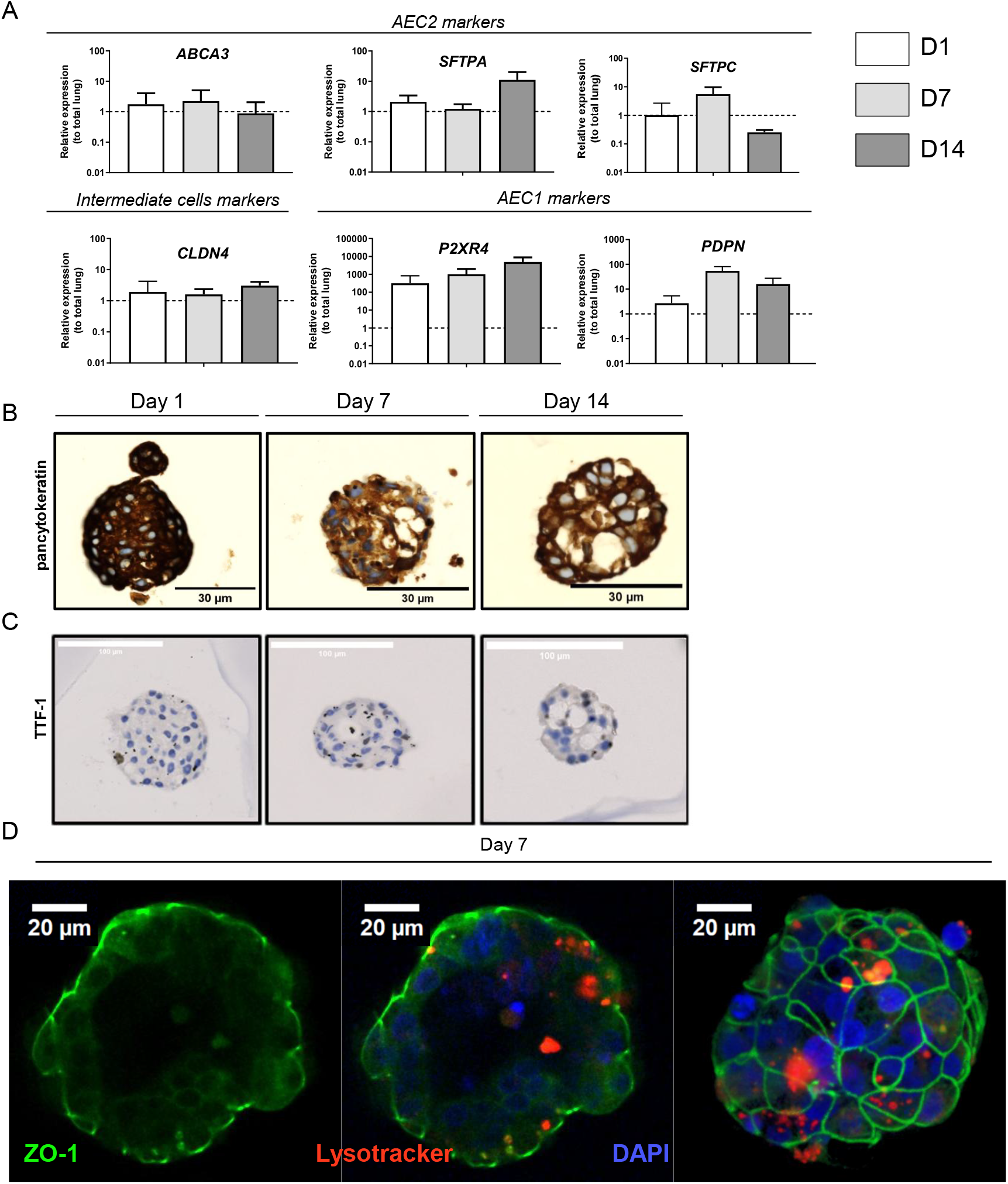
Characterization of cellular differentiation in the 3D-alveolospheres model. (A) qPCR results showed the persistence of AEC2 gene expression (*abca3, sftpa, sftpc*) over time, while markers of intermediate cells (*cldn4*) and AEC1 cells (*p2×4r, pdpn*) increased progressively n = 3 different donors (B) Bright-field images of H&E and pancytokeratin. One paraffin representative slice was assessed per donor per staining (C) Bright-field images of H&E and TTF1. One paraffin representative slice was assessed per donor per staining (D) fluorescent image of organoids stained for epithelial junction (ZO-1, green) and AEC2s (lysotracker, red) and nuclei (DAPI, blue) showing epithelial barrier AEC2s at day 1 AEC2: alveolar type 2 cell, *abca3: ATP Binding cassette Subfamily A Member 3, cldn4: claudin 4*, HTII-280: Human type 2 cells-280kDa protein, *p2×4r: purinergic receptor 2×4, pdpn: podoplanin, sftpa: surfactant protein A, sftpc: surfactant protein C, TTF-1: Thyroid Transcription Factor 1, ZO-1: Zona Occludens*

### Electron microscopy characterization

Electron microscopy analyses revealed the presence of abundant mitochondria and numerous lamellar bodies, which represent a characteristic feature of mature AEC2 ((**Figure 4 A**). Notably, LBs’ mean diameter was around 600nm as previously described [20]. Besides, we identified different maturation stages of LBs: intracellular, extracellular coiled and uncoiled LB but also numerous lipid bodies (**Figure 4 A**), which suggests surfactant production and secretion. Additionally, we identified isolated LBs, connected LBs and LBs connected to the membrane as described (**Figure 4 A**) [20]. To precisely assess the presence and number of LB over time, we developed an AI-based tool able to detect and quantify of these organelles and applied the tool on whole TEM images, which had been previously segmented to delineate the cell membranes (**Figure 4 B and Figure S2 C**) and then reconstructed by SBF-TEM into 3D images (**Figure S2 D**). We found that the distribution of the number of LBs increased over time (**Figure 4 C**), similarly to the number of LBs per LBs-containing cells (**Figure 4 D**). Finally, the total number of detected cells per alveolosphere increased over time (**Figure 4 E**). Meanwhile, the number of LBs-containing cells (e.g. AEC2), relatively to cells without lamellar bodies (e.g. intermediate or AEC1) tended to decrease over time (**Figure 4 F**). Collectively, these data suggest the progressive differentiation of AEC2 into intermediate cells and possibly AEC1 into the alveolospheres.

**Figure 4:**
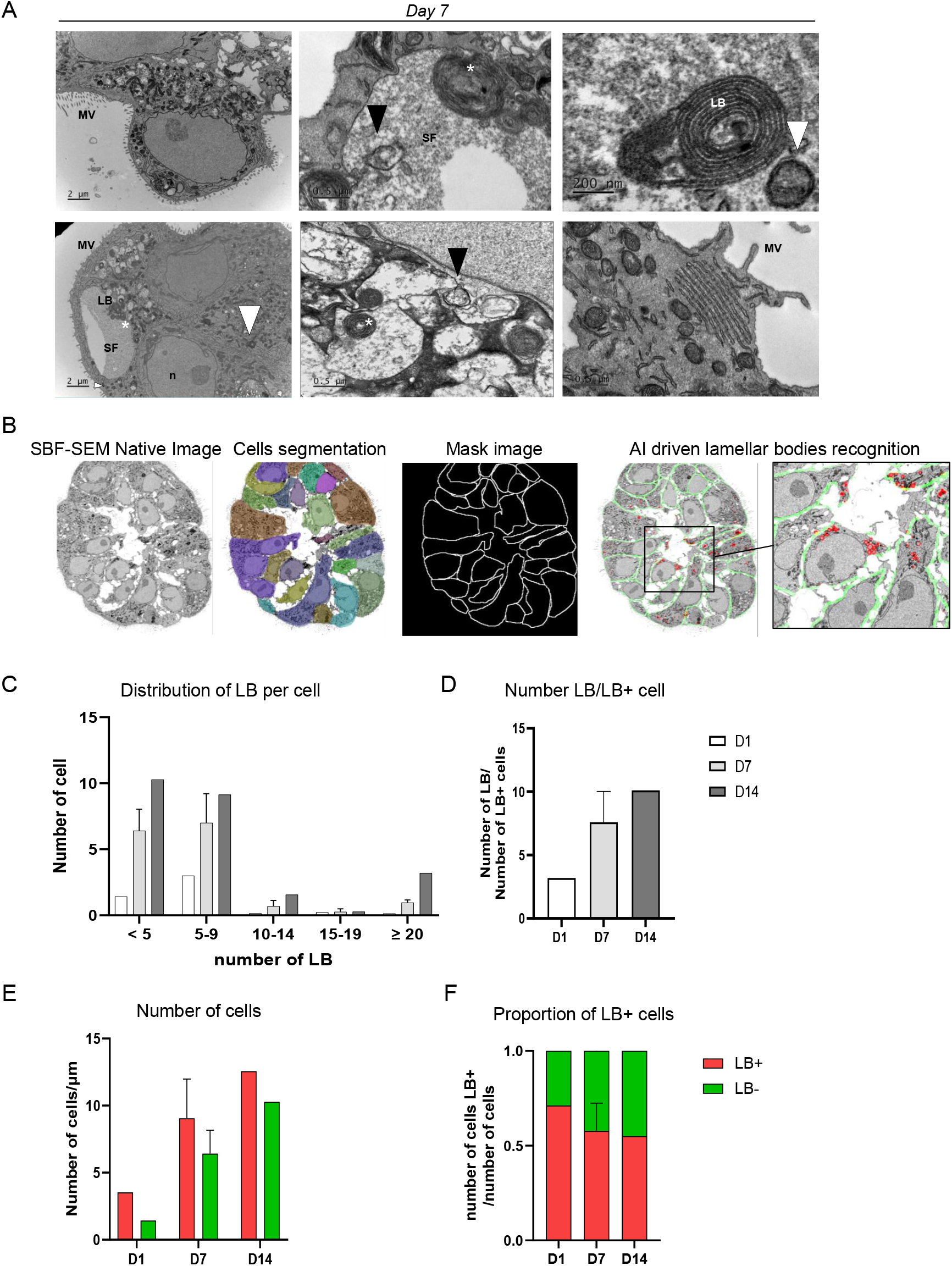
Characterization of cellular differentiation in the 3D-alveolospheres model by electron microscopy. (A) electron microscopic image of embedded organoid at day 7. LB indicate intracellular lamellar bodies, SF indicate surfactant, MV indicate microvillosities, n indicate nucleus. Extracellular lamellar bodies (white asterisks), uncoiled extracellular lamellar body (black arrow), lipid bodies (white arrows) (B) Reconstruction of an entire organoid with serial bloc face scanning electron microscopy (SBF-SEM), automatic cellular segmentation, and automatic detection of lamellar bodies in red Quantification was available for n=1 for D1, n=3 for D7, n=1 for D14) (C) Distribution of lamellar bodies (LB) per cell in organoids at day 1, 7 and 14 (D) Number of lamellar bodies (LB) per LB containing cells in organoids at day 1, 7 and 14 (E) Quantification the number of cells with (red) and without (green) lamellar bodies (LB) in organoids at day 1, 7 and 14 (E) Proportion lamellar bodies (LB) containing cells in organoids at day 1, 7 and 14 AEC2: alveolar type 2 cell, *AI: artificial intelligence, LB: lamellar body, MV: microvillosities, n: nucleus, pdpn: podoplanin, SBF-SEM:* serial bloc face scanning electron microscopy, *SF: surfactant*

Taken together, our 3D alveolosphere model meets criteria that identify organoids based on the definition previously published [4] : (i) composition of multiple organ-specific cell types, (ii) recapitulation of specific organ features such as lumen formation and barrier function, and, importantly, (iii) presence of lamellar bodies-containing AEC2 cells and differentiated intermediate cells.

### Emphysema modelling

We next assessed the ability of alveolospheres to accurately model emphysema. To this aim, we exposed alveolospheres to 1 or 5 % CSE, during 5 consecutive days (**Figure 5 A**). Global proteomic analyses of differentially expressed proteins between vehicle, 1 and 5% CSE-exposed alveolopsheres. We identified respectively 8502 and 554 up-regulated proteins (-Log2 fold change > 1.3) and 7501 and 1287 down-regulated proteins (Log2 fold change < -1.3). Five up-regulated (e.g. fatty acid metabolism, fibrogenesis and cell death) and ten down-regulated major pathways involved in emphysema (e.g. cell survival, immune response of cells) were identified after 5% CSE exposure (**Figure S7A)**. Exposure to CSE lead to the modification of pathways known to be involved in emphysema physiopathology. First, cell death through apoptosis was increased particularly with 5% CSE (**Figure 5 B, C, F)**. Second, cell-to-cell junctions are significantly disrupted as demonstrated with the modification of CEACAM1, integrin2 or SLP1 (secretory leukocyte protease inhibitor), one of the major defenses against the destruction of pulmonary tissues by elastase (**Figure 5 B, C)**. These results were confirmed through the top enriched canonical pathways where pulmonary healing signaling and extra-cellular matrix organization pathways were largely down-regulated after CSE exposure (**Figure 5 D, E, Figure S7B)**, similarly to sphere formation (**Figure 5 G)**. In addition, exposure to CSE also significantly increased inflammation such as TNF receptor or APOE **Figure 5 B, C)**, which is known to activate the NLPR3 inflammasome in BALF macrophages in lung diseases [21]. We identified between the top-enriched pathways activation virus pathogenesis and macrophages activation (**Figure 5 D, E, Figure S7C**) particularly after 5% CSE exposure, and 33 signaling (**Figure 5 H)**. Besides, CSE exposure also increased oxidative stress markers as shown via PDIA4 protein disulfide isomerase 4, calpastatin or CYP1A1 up-regulation (**Figure 5 B, C)**, also known to be up-regulated in COPD AEC2 (www.copdcellatlas.com) and ROS production (**Figure 5 I)**. In addition, proteins associated with stemness or precursor state are also up-regulated after CSE exposure such as SGB1A1, KRT4, TMEM45a and Ribosomal protein L22 like 1 **Figure 5 B, C)** are also over-expressed after CSE exposure, similarly as previous data particularly in organoids from COPD patients [22]. Thus, both considering that 5% CSE exposure seem to have more significant effect and previously published data [23], we chose to use 5% CSE exposure for the remaining experiments. CSE-exposed alveolospheres were smaller than control (**Figure 6 B, C**). Calcein staining was weaker in 5% CSE exposed-alveolospheres than in controls, suggesting altered cellular viability (**Figure 6 D**). Furthermore, the architecture appeared to be disrupted with the absence of central lumen within the alveolospheres exposed to 5% CSE (**Figure 6 E**). Epithelial barrier was also disturbed by CSE (**Figure 6 F**). Lamellar bodies were disorganized, large and clustered (**Figure 6 G**). Similarly, to what we found in the proteomic analyses, secreted inflammatory cytokines (MIF, IL-8 and PAI1) from CSE exposed-alveolospheres derived from 2 patients increased, according to the cytokine array (**Figure 6 H**) and we confirmed and increased in MIF concentration by ELISA in the supernatant of alveolospheres after 5% CSE (**Figure 6 I**). Finally, the 5% CSE exposure also increased oxidative stress gene markers assessed by RT-qPCR (*HMOX1, NQO1*, **Figure 6 J)**.

**Figure 5:**
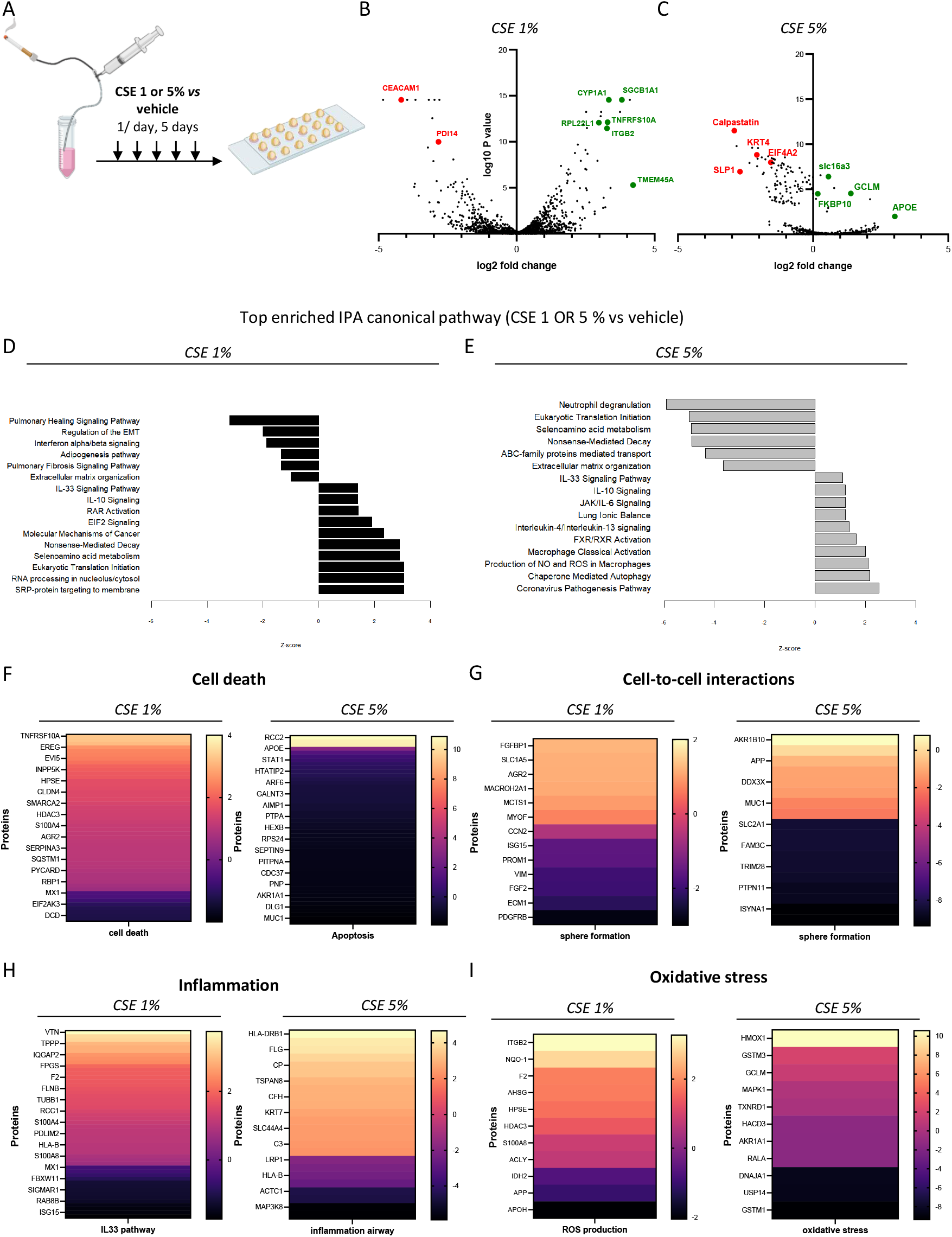
Emphysema modelling in the 3D-alveolosphere model by exposure to cigarette smoke extract (CSE) A. Alveolospheres were exposed daily for 5 consecutive days to 1 or 5% CSE. B. Volcano plot shows differential protein expression between 1% CSE and vehicle, from 3 donors. C. Volcano plot shows differential protein expression between 5% CSE and vehicle, from 4 donors. D Bar graphs illustrating the 16 top enriched Ingenuity Pathway Analysis (IPA) canonical pathways derived from differentially expressed genes of 1% (grey) CSE to vehicle, from 3 donors. E Bar graphs illustrating the 16 top enriched Ingenuity Pathway Analysis (IPA) canonical pathways derived from differentially expressed genes of 5% (black) CSE to vehicle, from 4 donors. F. Heatmap of cell death differentially expressed proteins between 1% CSE and vehicle exposed alveolospheres; and between 5% CSE and vehicle exposed alveolospheres(orange: activation; purple: inhibition). G. Heatmap of cell to cell interaction differentially expressed proteins between 1% CSE and vehicle exposed alveolospheres; and between 5% CSE and vehicle exposed alveolospheres(orange: activation; purple: inhibition). H. Heatmap of inflammation differentially expressed proteins between 1% CSE and vehicle exposed alveolospheres; and between 5% CSE and vehicle exposed alveolospheres(orange: activation; purple: inhibition). I. Heatmap of oxidative stress differentially expressed proteins between 1% CSE and vehicle exposed alveolospheres; and between 5% CSE and vehicle exposed alveolospheres(orange: activation; purple: inhibition). *CSE: cigarette smoke extract, IPA: Ingenuity Pathway Activity*

**Figure 6:**
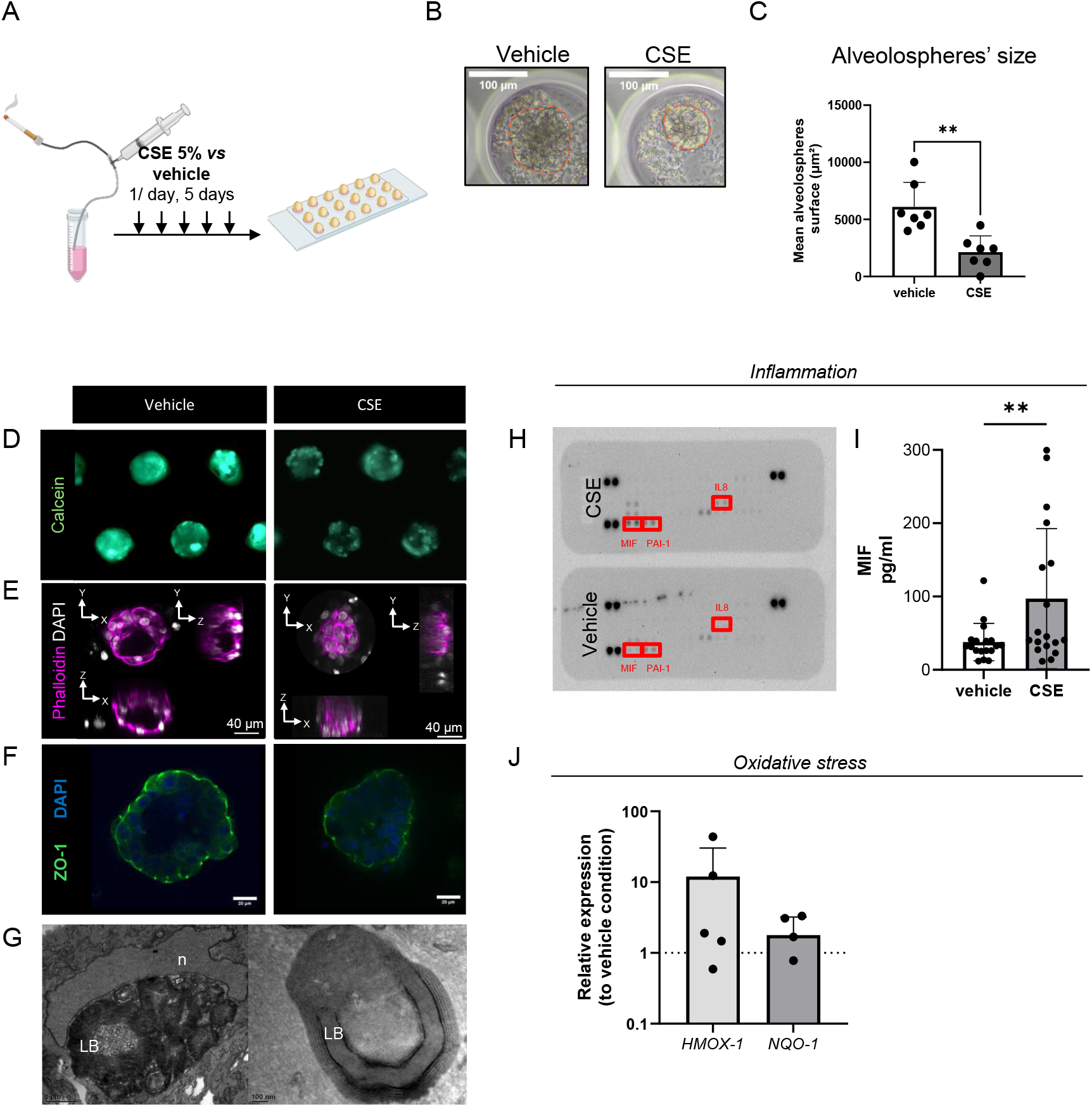
Emphysema modelling in the 3D-alveolosphere model by exposure to 5% cigarette smoke extract (CSE) A. Alveolospheres were exposed daily for 5 consecutive days to 5% CSE. B. Representative image of alveolosphere exposed to CSE *versus* vehicle C. Quantification of organoids surface exposed to 5% CSE or vehicle D. Viability, assessed by calcein staining was lower in 5% CSE exposed-alveolospheres than in vehicle exposed alveolospheres E. Self-organization appeared to be disrupted with the absence of central lumen within the alveolospheres exposed to 5% CSE, phalloidin in magenta F. 3D reconstruction organoid staining for Zona Occludens-1 (ZO-1, green); fluorescence and DAPI (blue) exposed to 5% CSE showing barrier dysfunction. G. Lamellar bodies are disorganized in 5% CSE-alveolospheres, LB: lamellar body, n: nucleus H. Induced inflammation in supernatant of alveolospheres exposed to 5% CSE compared to vehicle, assessed by cytokine array, showing an increase of MIF, PAI-1 and IL8 (n=2) I. ELISA for MIF confirmed an increased level of MIF in the supernatant of alveolospheres exposed to 5% CSE JInduced oxidative stress in alveolospheres exposed to 5% CSE compared to vehicle, assessed by qPCR (*HMOX-1, NQO-1*) *CSE: cigarette smoke extract*, DAPI: 4’,6-diamidino-2-phenylindole, *HMOX-1: heme-oxygenase-1, IL-8: interleukin-8, MIF: migration inhibitor factory, NQO-1: NAD(P)H dehydrogenase quinone 1, PAI-1: plasminogen activator inhibitor 1, ZO-1: Zona Occludens-1*

In summary, our data revealed that daily exposure to CSE of our standardized 3D-alveolospheres accurately recapitulate the induction of emphysema features, such as cell junction alteration, surfactant dysfunction, inflammation and oxidative stress.

## Discussion

Pathogenesis of emphysema involves impaired cellular junctions, surfactant dysregulation [7], chronic inflammation [1, 24] and oxidative stress in lungs. Many cellular and molecular processes are challenging to study *in vivo*, and traditional 2D *in vitro* cultures fail to replicate key lung features. Consequently, *in vitro* 3D culture systems have emerged as a valuable platform for better understanding pathological processes. Here, we developed a tunable, sustainable, and reproducible 3D-alveolosphere model derived from human primary AEC2 to investigate emphysema mechanisms. Using hydrogel calibrated microwells, we observed faithful replication of native lung characteristics, including lumen formation, epithelial barrier and epithelial cell composition of the human distal alveolar-like structures (AEC1 and AEC2). In addition, chronic CSE exposure leads to key pathophysiological features of emphysema such as increased cell death, cell junction alteration, oxidative stress and inflammation. Finally, we managed to develop an innovative AI-driven tool to characterize alveolospheres maturation via SBF-TEM.

Our alveolar organoid model presents several strengths. Until now, traditional methods for alveolosphere culture suffer from a low reproducibility and geometric heterogeneity [8, 25] limiting the scope and application of organoid research in the field of respiratory diseases. Here, AEC2 seeded into microwells of spherical shape self-organized into spherical structures, with the progressive formation of a central lumen. While luminogenesis mechanisms remain unclear, curved geometry [26], fluid pressure [27] and actin polymerization followed by both ion flux to intracellular space and tight junction formation [28] have been identified as key factors. It has also been shown that an apical actin network including tropomyosin, cytokeratin-based intermediate filaments, increases in size as lumen grows [29]. Then, tight junction formation occurred, allowing lumen growth by the establishment of an osmotic pressure gradient [28]. Our alveolospheres model, by generating large numbers of uniform distal lung organoids, could provide a high-throughput method to study luminogenesis mechanisms more accurately.

Second, it is adjustable, enabling us to model alveolospheres whose 200µm diameter is close to existing alveoli *in vivo* [18]. Also, the hydrogel matrix stiffness can be modulated to obtain the measurements similar to those described in emphysema (∼1kPa) compared to native control lung (∼ 5 kPa) [13–15]. The impact of matrix stiffness in lung diseases is of growing interest in emphysema with decrease in elastic recoil [1]. At the microscale level, differentiated AEC1s, originating from AEC2s, both of which are sensitive to mechanical stress and rigidity. Aberrant mechanical stretching and modification of stiffness of the extracellular matrix, such as observed in emphysema, can result in cellular barrier dysfunction, metabolic dysfunction, cytotoxicity, and inflammation. In emphysema, the mechanical forces of normal breathing [30] cause inflammation-related [31] or elastase-induced [32] remodeling of the septal wall, weakening the collagen, elastin, and the entire alveolar wall [33, 34]. Interestingly, de Hilster *et al*. tried to develop an *in vivo* hydrogel directly derived from human lung patients, decellularized, reduced to a powder, and further reconstituted. However, they failed to reproduce low emphysema stiffness [35]. Overall, there are still many unanswered questions regarding the micromechanics of the alveolus, especially in the context of emphysema. Therefore, our tunable model may be of interest particularly to mimic this important property in emphysematous lungs.

Many studies have shown that the growth of 3D-cultured alveoli may be dependent on co-cultured fibroblasts, suggesting the essential role of fibroblast-derived niche factors in the maintenance of alveolar epithelium [25, 36]. In a recent murine study, Yao *et al*. were able to demonstrate the critical role of fibroblasts to mediate alveolar regeneration [37], in air-liquid interface cultures. In the current study, we were able to maintain alveolospheres in culture during 14 days without a fibroblast niche, through appropriate medium and hydrogels substrates. Therefore, again through the specific properties of our hydrogel and available commercial media, we describe a robust method for isolation and 3D culture of human AEC2 to obtain standardized alveolospheres that exhibit distinct alveolar-like lung structures until 14 days of culture. Future studies will aim at determining whether incorporation of fibroblasts in our model promotes long-term maintenance and further differentiation.

Long-term maintenance of alveolar organoids derived from human adult differentiated stem cells with differentiated AECs, particularly AEC1, is critically missing. Single cell RNA sequencing has enabled the identification of a previously unknown intermediate cell state during the differentiation of AEC2 into AEC1, particularly after lung injury. Several molecular markers have been identified in this transient state, including krt-8 and claudin-4, while expression of AEC1 and AEC2 markers is low [38, 39]. Here, our results suggest the presence of these intermediate cells, evidenced by the expression of claudin-4, highlighting the existence of a dynamic differentiation process. In addition, using AI-driven analysis of TEM data, we can also suspect the presence of AEC1, negative for LB presence, highlighting the robustness of this standardized tool leading to key features of mature *in vivo* alveoli. Moreover, we have also demonstrated the increase of proteins that are classically expressed in AEC2 progenitor cells as described in the literature such as SGB1A1 and TMEM45a, particularly involved in COPD patients [22, 39–41]. Therefore, our data also indicate that these 3D culture models offer a valuable approach for studying emphysema mechanisms. It is well known that repeated aggressions, such as exposure to cigarette smoke, lead to chronic inflammation and oxidative stress both *in vivo* and *in vitro* [42, 43]. Chan *et al*. have shown the establishment and characterization of COPD lung organoids from nasopharyngeal and bronchial human cells. They were able to recapitulate the *in vivo* existing differences between non-diseased and COPD organoids. However, there were no data at the alveolar level from emphysematous patients. Notably, SCGB1A1 was recently identified in the AEC2 of alveoli particularly in COPD lungs [22, 40, 44]. Interestingly, we also found an over expression of TMEM45a which is overexpressed in so-called rbAEC2, which have been described as progenitor cells and increased in emphysematous lung [22]. In addition, inflammation process after chronic CSE exposure led to MIF secretion in our alveolospheres, as previously shown in mouse model of COPD secondary to tobacco [45]. Therefore, we think that our model, capable to reproduce classical pathways of emphysema after chronic cigarette exposure, will allow larger experiments.

The study has several limitations that also need to be discussed. First, the alveolospheres do not fulfill the whole microwells however, the mean size remains stable and reproducible over time. Second, while the presence of LB at different stages of maturation suggests surfactant secretion into the alveolosphere lumen, the specific components of this lumen could not be explored or directly exposed to CSE in our model. Indeed, accessing the apical surface remains challenging because it is enclosed within the alveolosphere. However, further experiments to reverse alveolosphere polarity would be feasible while maintaining proper polarity and barrier function as previously described in enteroids by modifying the substrate’s stiffness [46]. We can also modulate the stiffness offered by our hydrogel microstructuration method [12].Finally, although the presence of microvilli and tight junctions are strong indicators of cell orientation, the polarity of cells remains to be clearly determined. Nevertheless, this model can provide a standardized platform for elucidating emphysema pathways and offers insights into potential therapeutic interventions.

## Conclusion

This study aimed at developing a standardized and reproducible 3D-model of emphysema using human AEC2 isolated from lung samples in hydrogel microwells of adjustable shape, size and stiffness. By overcoming the limitations of previous models, such as heterogeneity in size and shape, this innovative approach allows for a more accurate representation of lung tissue *in vitro*. Results indicated that the 3D-alveolospheres could be maintained in culture for 14 days, with progressive formation of a central lumen resembling *in vivo* alveolar structures. Tight junctions and epithelial barrier formation were observed, along with expression of AEC1 and AEC2 markers and confirmation of surfactant synthesis through TEM images. Chronic exposure to CSE resulted in decreased cell viability, architectural disorganization, and increased oxidative stress and inflammatory markers, accurately reproducing emphysema pathophysiology, providing a valuable tool for investigating mechanisms involved in emphysema, particularly through exposure to CSE.

## Declarations

### Consent for publication

All authors have read and approved the manuscript.

## Acknowledgements

We thank the study participants and the staff of the Thoracic Surgery, Respiratory, Lung Function Testing departments from the University Hospital of Bordeaux (Bordeaux, France); Isabelle Goasdoue, Isabelle Bernis, Natacha Robert, Marine Douillet, and Rachel Gilson from the Clinical Investigation Center for technical assistance; Magali Mondin and Sébastien Marais for help with imaging and image analysis at the Bordeaux Imaging Centre (BIC; Bordeaux, France) and Atika Zouine, Vincent Pitard and Jean-Michel Griffon for technical assistance at the Flow cytometry facility, UAR CNRS 3427, INSERM US 05, Univ. Bordeaux, F-33000 Bordeaux, France (CNRS UMS 3427, INSERM US 005, Univ. Bordeaux, F-33000 Bordeaux, France). Microscopy was performed at the BIC, a service unit of the CNRS-INSERM, and Bordeaux University, a member of the National BioImaging Infrastructure of France supported by the French National Research Agency (ANR-10-INBS-04). We thank Pr Sylvain Gabriele and Lucie Ergot (Université de Mons) for performing stiffness measurements of hydrogels using Optics 11 Chiaro.

## Availability of data and material

The datasets used and analyzed during the current study are available from the corresponding author on reasonable request.

## Competing interests

MG CJ YB MT ES KR EL VS have nothing to disclose

AP reports grants and personal fees from Chiesi, GSK

HB GM FD JWD AAR PE PB LG EM GD

ID has a patent (EP N°3050574).

PH reports grants and non-financial support from Avad, non-financial support from Chiesi and GlaxoSmithKline, outside the submitted work.

MZ reports grants and personal fees from AVAD, Boehringer Ingelheim, personal fees from Chiesi, personal fees from Sanofi, personal fees from CSLBehring, personal fees from AstraZeneca, personal fees from Menarini and personal fees from GSK, and support for attending meetings and/or travel from Chiesi, AstraZeneca and GSK outside the submitted work

