## Supplementary material for "Tunable 3D Alveolosphere Model from Human Alveolar Cells: A Breakthrough Tool to Explore Emphysema Pathophysiology": TABLES

| | N= 52 |
|---|---|
| **Age**, years | 65 [58 ; 71] |
| **Gender**, female male | 27 (52) |
| **BMI,** kg/m$^2$ | 25.3 [22.5 ; 27.0] |
| **Smoking,** pack-year | 29 [10 ; 50] |
| **Smoking habits** | |
| Never smoker | 9 (17) |
| Former smoker | 36 (69) |
| Current smoker | 7 (13) |
| **FEV1/FVC<0.7** | 6 (12) [a] |
| **FEV1/FVC** | 0.73 [0.72; 0.82] [a] |
| **FEV1**, L | 2.21 [1.80; 2.71] [a] |
| **FEV1**, % predicted | 86 [73; 108] [a] |
| **FVC**, L | 2.98 [2.28; 3.63] [a] |
| **RV,** L | 2.46 [1.78; 2.82] [b] |
| **RV,** % predicted | 119 [90; 130] [b] |
| **TLC,** L | 6.46 [4.73; 6.57] [b] |
| **TLC,** % predicted | 100 [86; 113] [b] |
| **DLCO,** % | 67 [54; 86] [c] |
| **$PaO_2$**, mmHg | 74 [71; 88] [d] |
| **pH** | 7.43 [7.41; 7.46] [d] |
| **Emphysema (LAA%<950 HU)** | 4 (8) [e] |
| **Cancer** | |
| Adenocarcinoma | 31 (60) |
| Squamous cell cancer | 5 (10) |
| Other | 7 (13) |
| **Transplantation** | |
| Emphysema | 4 (8) |
| Other | 5 (10) |
| **Lung samples** | |
| Weight (grams) | 2.8 [1.4; 3.9] [f] |
| 2D culture (days) | 25 [20; 27] [g] |
| Total number of cells (.10$^6$) | 308 [107; 401] [h] |
| Number of HTII-280+ cells (.10$^6$) | 3.9 [1.15; 3.95] [i] |

**Table 1**: Patients' characteristics (2D = 2 dimension, BMI = body mass index, DLCO= diffusing capacity of lung for carbon monoxide, FEV1 = forced expiratory volume in 1 second, FVC = forced vital capacity, HU = Hounsfield Units, $PaO_2$ = arterial oxygen pressure, RV = residual volume, TLC = total lung capacity). Low attenuation area (LAA%) was defined as the percent of voxels below −950 HU. Data are presented in n (%) or median [quartile 1; quartile 3]

Missing values are [a] n=3, [b] n=6, [c] n=15, [d] n=28, [e] n=13, [f] n=4, [g] n=3, [h] n=8, [i] n=4
