## SUPPLEMENTARY TABLE for "Tunable 3D Alveolosphere Model from Human Alveolar Cells: A Breakthrough Tool to Explore Emphysema Pathophysiology"

**SUPPLEMENTARY MATERIALS**

| Gene | Target | forward | reverse |
|---|---|---|---|
| *gus b*<br>Glucuronidase β | Housekeeping gene | 5'-CCATCTGGGTCTGGATCAAAA | 5'-TGAAATCGGCAAAATTCCAAAT |
| *p2xr4*<br>Purigenic receptor P2X4 | AEC1 | 5'-TGGCCGATTATGTGATACCAGC | 5'-CACACAGTGGTCGCATCTGGAA |
| *pdpn*<br>Podoplanin | AEC1 | 5'-TGCCGAAGATGATGTGGTGAC | 5'-GGACTGTGCTTTCTGAAGTTGGC |
| *abca3*<br>ATP binding cassette subfamily A member 3 | AEC2 | 5'-TTGACAGTCGCAGAGCACCTT | 5'-CTCCGTGAGTTCCACTTGTCCT |
| *sftpa*<br>Surfactant protein A | AEC2 | 5'-GGCACCCTTAGCCACTTCAT | 5'-GAGGGCTCCATCTCATGTCG |
| *sftpc*<br>Surfactant protein C | AEC2 | 5'-CTCTCTGCAGGCCAAGCCCG | 5'-TTCCACTGACCCGGAGGCGT |
| *cldn4*<br>Claudin 4 | AEC1 | 5'-GGCTGCTTTGCTGCAACTGTC | 5'-GAGCCGTGGCACGTTACACG |
| *cxcl8*<br>Interleukin 8 | Inflammation | 5'-AAGGCGGCCAGGATATAACT | 5'-TTGGGCCAACAGTAGCCTTC |
| *hmox*<br>Heme oxygenase | Oxydative stress | 5'-GAATCGAGCAGAACCAGCCT | 5'-CTCAGCATTCTCGGCTTGGA |
| *nqo1*<br>NADPH quinone deshydrogenase1 | Oxydative stress | 5'-TAGCCTGTAGCCAGCCCTAA | 5'-GCCTCCTTCATGGCGTAGTT |
| *srxn1*<br>Sulfiredoxin-1 | Oxydative stress | 5'-AGCACATTAGCAGGTCAACCA | 5'-AGGCTATTGCATGGTGTGT |

**Suppl Table 1:** Summary of primers used for qPCR
AEC1 = alveolar epithelial cell type1, AEC2 = alveolar epithelial cell type2

| **Antibodies** | **Target** | **Reference** | **Dilution** |
|---|---|---|---|
| IgM anti HTII-280 | AEC2 | TB-27A HT2-280 | 1:100 |
| Anti IgM A488 or A647 | Secondary | A-21042 ou A-21238 | 1:500 |
| DAPI | Nucleus | D1306 | 1:10000 |
| Phalloidin | Actin | A-22284 | 1:40 |
| ZO-1 | Tight Junction | 339100 | 1:100 |
| Anti IgG A568 | Secondary | A-11004 | 1:500 |
| Lysotracker™ Red Dnd-99 | Lamellar bodies | L7528 | 50 nM |
| Pancytokeratin | Epithelial cells | sc8018, Santa Cruz | 1 :25 |
| CD45 | Leucocytes | 555485, BD Pharmingen | 1 :25 |

**Suppl Table 2**: Summary of antibodies (and associated references) for immunolabelling. AEC2 = alveolar epithelial cell type2, DAPI = 4'6-diamidino-2-phenylindole, ZO-1 = zonula occludens protein 1.

| | **N** | **%** |
|---|---|---|
| **Medications** | | |
| LABA or LAMA | 6 | 11.5 |
| LABA and LAMA | 5 | 9.6 |
| LABA and ICS | 0 | 0 |
| LABA and LAMA and ICS | 2 | 3.8 |

**Suppl Table 3**: Supplemental patients' characteristics (LABA: long acting beta-2-agonists, LAMA: long acting anti-muscarinic, ICS: inhaled corticosteroids)

| | |
|---|---|
| **Professional, technical, and related workers** | |
| Medical, dental, veterinarians, pharmacists | 1 |
| Nurses, midwives, medical x ray technicians | 3 |
| Architects, engineers, and related technicians | 1 |
| Statisticians, mathematicians, systems analysts | |
| Teachers | 2 |
| Social workers | |
| Chemists, biologists, and related workers | 1 |
| Jurists, journalists, and related workers | |
| Other professional and technical workers | 1 |
| **Managerial workers** | 2 |
| **Clerical and related workers** | |
| Stenographers, typists | 3 |
| Bookkeepers, cashiers | |
| Other clerical and related workers | |
| **Sales workers** | |
| Technical saleswomen, commercial travelers | 1 |
| Saleswomen, shop assistants | 1 |
| Other sales workers | 1 |
| **Service workers** | |
| Cleaners and helpers | 3 |
| Nurses' aides | 2 |
| Hairdressers, beauticians | 1 |
| Other service workers | 2 |
| **Agricultural workers** | 1 |
| **Production workers** | |
| Textile workers | |
| Food and beverage processors | 2 |
| Electronic or metal processors | 1 |
| Material handlers and related equipment operators | 3 |
| Other production workers | 2 |
| **Housewife** | 2 |
| **Other or unknown occupations** | 16 |

**Suppl Table** 4: Description of occupational activities among the cohort

| | Final analyses | 2D failure |
|---|---|---|
| | N= 52 | N=14 |
| **Age**, years | 65 [58 ; 71] | 66 [61 ; 70] |
| **Gender**, female | 27 (52) | 7 (54) |
| **BMI,** kg/m$^2$ | 25.3 [22.5 ; 27.0] | 24.9 [22.9 ; 27.2] |
| **Smoking,** pack-year | 29 [10 ; 50] | 41 [27.5 ; 57.5] |
| **Smoking habits** | | |
| Never smoker | 9 (17) | 3 (23) |
| Ex-smoker | 36 (69) | 5 (38) |
| Current smoker | 7 (13) | 4 (31) |
| Missing data | 0 | 1 |
| **FEV1/FVC<0.7** | 6 (12) [a] | 11 (85) * |
| **FEV1/FVC** | 0.73 [0.72; 0.82] [a] | 0.61 [0.51; 0.68] * |
| **FEV1**, L | 2.21 [1.80; 2.71] [a] | 1.83 [1.5; 2.46] * |
| **FEV1**, % predicted | 86 [73; 108] [a] | 69 [49; 96] * |
| **FVC**, L | 2.98 [2.28; 3.63] [a] | 2.87 [2.50; 3.67] |
| **RV,** L | 2.46 [1.78; 2.82] [b] | 3.69 [2.50; 3.96] * |
| **RV,** % predicted | 119 [90; 130] [b] | 170 [112; 187] * |
| **TLC,** L | 6.46 [4.73; 6.57] [b] | 6.64 [5.07; 7.58] * |
| **TLC,** % predicted | 100 [86; 113] [b] | 113 [99; 116] * |
| **DLCO,** % | 67 [54; 86] [c] | 65 [59 ;74] [j] |
| **$PaO_2$**, mmHg | 74 [71; 88] [d] | 82 [62; 96] [k] |
| **pH** | 7.43 [7.41; 7.46] [d] | 7.46 [7.44; 7.48] |
| **Emphysema (LAA%< 950 HU)** | 4 (8) [e] | 0 [l] |
| **Cancer** | | |
| Adenocarcinoma | 31 (60) | 9 (69) |
| Squamous cell cancer | 5 (10) | 0 |
| Other | 7 (13) | 1 (8) |
| **Transplantation** | | |
| Emphysema | 4 (8) | 2 (9) |
| Other | 5 (10) | 1 (27) * |
| **Lung samples** | | |
| Weight (grams) | 2.8 [1.4; 3.9] [f] | 1.95[0.7; 3.15] [m] |
| 2D culture (days) | 25 [20; 27] [g] | 0 |
| Total number of cells (.10$^6$) | 308 [107; 401] [h] | 212 [74; 356] [m] |
| Number of HTII-280+ cells (.10$^6$) | 3.90 [1.15; 3.95] [i] | 6.10 [0.9; 4.97] [m] |

**Suppl table 5:** Comparison of the characteristics of patients whose samples were included in 3D analyses *versus* those whose samples failed in 2D cell culture.

Data are presented in n (%) or median [quartile 1; quartile 3];

2D = 2 dimension, BMI = body mass index, DLCO= diffusing capacity of lung for carbon monoxide, FEV1 = forced expiratory volume in 1 second, FVC = forced vital capacity, HU = Hounsfield Units, $PaO_2$ = arterial oxygen pressure, RV = residual volume, TLC = total lung capacity). Low attenuation area (LAA%) was defined as the percent of voxels below −950 HU. * $p<0.05$.

Missing values are [a] n=3, [b] n=6, [c] n=15, [d] n=28, [e] n=13, [f] n=4, [g] n=3, [h] n=8, [i] n=4, [j] n=4, [k] n=3, [l] n=11, [m] n=3
