## SUPPLEMENTARY MATERIAL for "Tunable 3D Alveolosphere Model from Human Alveolar Cells: A Breakthrough Tool to Explore Emphysema Pathophysiology"

**Supplementary material - Innovative Tunable 3D Alveolosphere Model from Human Alveolar Cells: A Breakthrough Tool for Exploring Emphysema Pathophysiology**

***Lung tissue processing***

Lungs were dissociated mechanically and enzymatically with type I collagenase (Ref 17100017, 500 UI/g at the dilution of 1:10 in Hanks' balanced salt solution (HBSS) with calcium, Thermofisher) for 45 minutes at 37°C. This homogenate was filtered through a 100µm then 70µm sieve to remove the biggest remaining debris.
Then, cells were labeled with an anti-HTII-280 monoclonal mouse IgM antibody (Terrace Biotech, San Francisco, CA). After washing, cell suspension was labelled with magnetic beads coupled to an anti-IgM antibody (Milteny, 130-047-301), enabling magnetic sorting (LS column, Milteny Biotech 130-042-401).

***Immunostaining***

The alveolospheres were fixed in 4% Paraformaldehyde Solution (PFA, ThermoFisher Scientific), permeabilized with BSA 1%/Triton X-100 0.1% for 30 minutes and saturated with PBS/1% bovine serum albumin (BSA, Sigma) for 1 hour at room temperature. Primary antibodies were incubated at 4°C overnight. After PBS washing, secondary antibodies coupled to fluorophores were added for 1 hour at room temperature (**suppl table 2**). Lamellar bodies were stained by lysotracker(L7528) according to manufacturer's datasheet. Images were acquired at Bordeaux Imaging Center (BIC) using a confocal spinning-disk microscope (Spinning-disk LiveSR Microscope, Leica DMi8 Yokogawa CSU-W1 + ILAS2).

***Transmission electron microscopy***

The samples were fixed with 2% PFA and 2.5% glutaraldehyde in 0.1M cacodylate buffer at pH 7 for 1 hour at room temperature (RT). After washing, the organoids were treated with osmium tetroxide (2% wt/vol) and potassium ferrocyanide (1.5% wt/vol) in 0.1 M cacodylate buffer for 60 min at 4°C, and then washed with ultrapure water. Samples were incubated for 20 min in a thiocarbohydrazide solution (1% w/v in water) at RT, then washed with ultrapure water and then incubated in 2% osmium tetroxide in water at RT for 30 min. After washing, organoids were incubated in 2% uranyl acetate at 4°C overnight. In order to increase the contrast, Walton's lead aspartate staining was performed for 30 min at 60°C. After final washing steps in ultrapure water, the samples were dehydrated in a graded ethanol series and acetone. After impregnation, organoids were embedded in epoxy resin and incubated at 60°C for 48 h.

For transmission electron microscopy, ultrathin sections of 70 nm thick were cut and picked up on copper grids. Samples were examined with a Transmission Electron Microscope (H7650, Hitachi, Tokyo, Japan) at 80 kV. For SBF-SEM, embedded samples were mounted on aluminium specimen pins using a silver filled conductive resin. Organoids were further trimmed with a diamond knife on an ultramicrotome (Leica EM UC7). SBF-SEM acquisition was performed with a ZEISS GeminiSEM300 with a fitted Gatan 3View2XP system (Gatan, Abingdon,

UK). The acquisition parameters were as follows: Pixel size 15 nm, Slice thickness 70 nm (Z slice), accelerating voltage 1.2 Kv and using a focal load compensation. Images were collected from electrons backscattered with an OnPoint Detector (On point - Gatan Inc., Pleasanton, CA, USA) using Digital Micrograph software (Gatan).

***Lamellar bodies (LB) quantification by artificial intelligence (AI) with transmission electron microscopy (TEM) serial block face (SBF) images***

The processing of electron microscopy images to classify organoid cells based on their lamellar body (LB) content was performed following a three-step protocol: training the Ilastik model, training the Cellpose model, and analyzing lamellar body density.

*Training the Ilastik Model*

To train the detection algorithm for LB, we used serial block-face scanning electron microscopy (SBF-SEM) images. The native images were first preprocessed to standardize contrasts across stacks acquired at different times. The resolution was reduced by 80%, and the contrast was adjusted to remove background noise and reveal intracellular structures. These preprocessing steps were performed using ImageJ.

Using the Ilastik software and its "pixel classification" feature, we created a model to automatically segment lamellar bodies within cells. For this, a training dataset was created using crops from various acquisitions of SBF-SEM images derived from preprocessed stacks. Crops were taken in three axes to enhance the model's robustness. The first step involved tagging the following structures: "Background," "Cell," and "Lamellar Bodies" on several crops. After analyzing the tags, the model proposed a generalized labeling for the entire image. The second step involved correcting the model's tagging errors, followed by reanalysis. These two steps were repeated until a satisfactory and generalizable result was achieved. The segmentation performed by the model was subsequently used (**figure S2 A, B**).

*Training the Cellpose Model*

To create a segmentation mask for organoid cells, we trained a second model using the Cellpose software. Reduced-resolution images were used (maximum resolution of 500x500 pixels). Orthogonal slices (XY, XZ, YZ) were extracted to form the training dataset.

Training the Cellpose model involves an initial manual segmentation of cells on the extracted slices. The generated masks were progressively refined through successive iterations. Once the segmentation was deemed satisfactory, the model was applied to the complete stack. The final mask obtained was saved for future use (**figure S2 C**), and we performed 3-D reconstruction of serial bloc face scanning electron microscopy (SBF-SEM) with artificial intelligence lamellar bodies recognition

*Analyzing Lamellar Body Density*

The final step involves analyzing the masks generated by Ilastik (lamellar bodies) and Cellpose (cells) in ImageJ. The volume of LB within each cell was measured using a Python script. Only cells with a volume above a predefined threshold (≥ 1000 voxels) were included in the final analyses. The analyses were then performed using GraphPad software. This protocol enables precise segmentation of structures of interest and robust quantification of lamellar bodies at the cellular level. Quantification has been assessed for n=1 at D1, n= 3 at D7 and n= 1 at D14.

***Flow cytometry***

Gates for all wavelengths were set by unstained cells and isotype controls for each antibody. Cells were analyzed by fluorescence activated cell sorting (FACS) Canto™ (BD, Biosciences, USA), and the results were analyzed using FACSDiva™ v9.0 (BD, Biosciences, USA). Gates for all wavelengths were set by unstained cells and isotype controls for each antibody. Fluorescence-minus-one (FMO) controls were used whenever possible for positive/negative population gating.

***Label-free quantitative proteomics***

The proteins were desalted and digested using the Single-pot, solid-phase-enhanced sample preparation (SP3) method (X2)[16]. NanoLC-MS/MS analysis were performed using a Vanquish Neo UHPLC System (Thermo Scientific) associated to Orbitrap Exploris™ 480. The peptide extracts were loaded onto a 5 mm × 300 µm ID PepMap Neo Trap Cartridge (C18, 5 µm particle size, 100 µ pore size, Thermo Scientific) and separated on an analytical column (25 cm × 75 µm ID, 1.7 µm, C18 beads, Ionopticks) at a flow rate of 300 nL/min at 50°C using a multistep gradient of 3–25% mobile phase B (80% MeCN in 0.1% formic acid) for 45 minutes and 25–35% B for 15 minutes, 35–95% B for 1 minute and an 11-minute wash at 99% B. The mass spectrometer operated in positive ion mode at a 1.4 kV needle voltage, and data were acquired using Xcalibur 4.5 software in a data-dependent mode. MS scans (m/z 375–1500) were recorded at a resolution of R = 120000 (@ m/z 200), a standard AGC target, and an injection time in automatic mode, followed by a top speed duty cycle of up to 1 second for MS/MS acquisition. Precursor ions (2–6 charge states) were isolated in the quadrupole with a mass window of 2 Th and fragmented with HCD @ 30% normalized collision energy. MS/MS data were acquired with a resolution of R = 15000 (@m/z 200), a standard AGC target, and a maximum injection time in automatic mode. Selected precursors were excluded for 45 seconds.

Protein identification and label-free quantification (LFQ) were done in Proteome Discoverer 3.1. The CHIMERYS node using the prediction model inferys_3.0.0 fragmentation was used to identify proteins in batch mode by searching against the UniProt Homo sapiens database (82233 entries, released June 2024). Two missed enzyme cleavages were allowed for trypsin. Peptide lengths of 7–30 amino acids, a maximum of 3 modifications, charges of 2–4, and 20 ppm for fragment mass tolerance were set. Oxidation (M) and carbamidomethyl (C) were respectively searched as dynamic and static modifications by the CHIMERYS software. Peptide validation was performed using the Percolator algorithm and only "high confidence" peptides were retained corresponding to a 1% false discovery rate at the peptide level. Minora feature detector node (LFQ) was used along with the feature mapper and precursor ions quantifier. The normalization parameters were selected as follows: (1) Unique peptides, (2) Precursor abundance based on intensity, (3) Normalization mode: total peptide amount, (4) Protein abundance calculation: summed abundances, (5) Protein ratio calculation: pairwise ratio based and (6) Missing values were replaced with random values sampled from the lower 5% of detected values. Quantitative data were considered for master proteins, quantified by a minimum of 2 unique peptides, a fold changes above 2 and a statistical p-value adjusted using Benjamini-Hochberg correction for the FDR lower than 0.05.

***Emphysema quantification on CT-scans***

CT were acquired at full inspiration in the supine position (GE Revolution®machine), with the following parameters: 120 kV, 250 mAs, 1-mm slice thickness, and standard reconstruction. CT doses ranged from 1 to 3 mSv. CT images using in-house software written in Matlab (Natick, Massachusetts: The MathWorks Inc.) as a surrogate for emphysema extension were analyzed by trained radiologists and pulmonologists (GD, AP), by using the density mask principle.
