## Supplementary material for "Tunable 3D Alveolosphere Model from Human Alveolar Cells: A Breakthrough Tool to Explore Emphysema Pathophysiology": SUPPL FIGURES

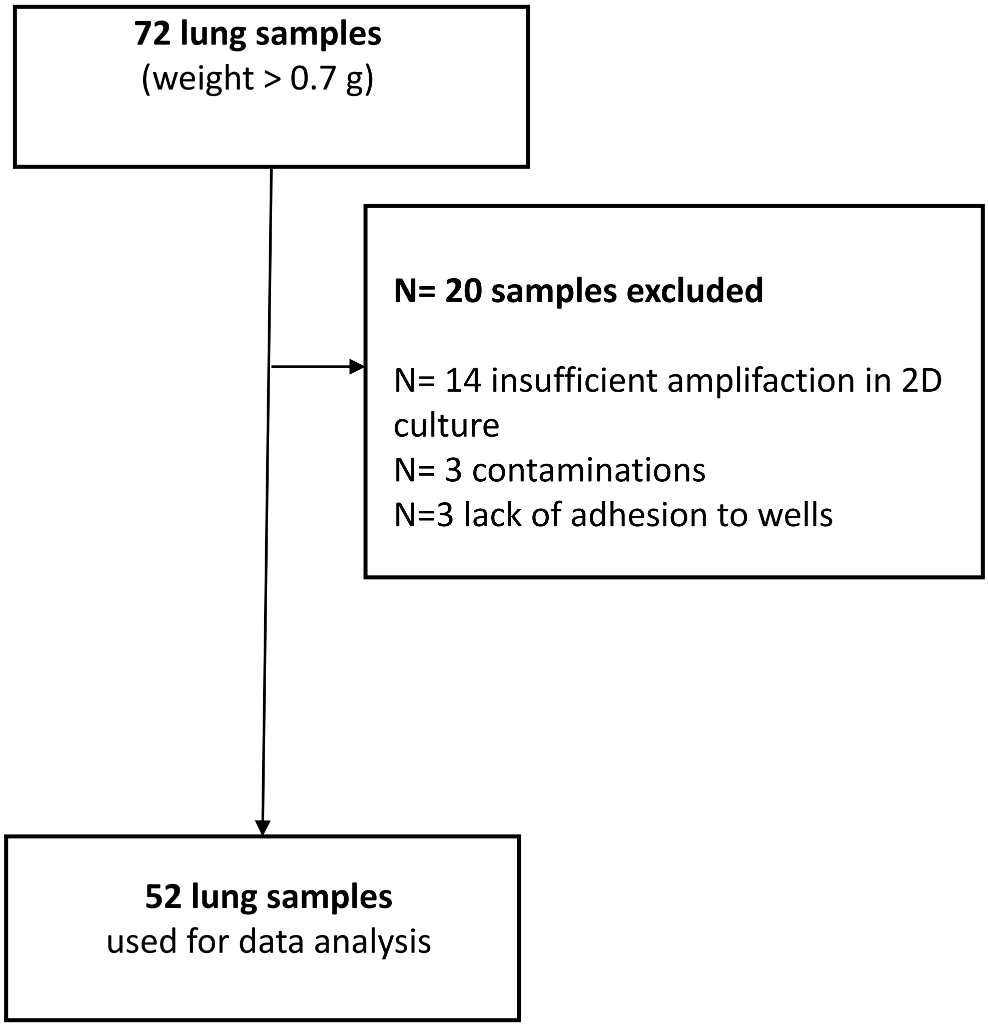
**72 lung samples**
(weight > 0.7 g)
**N= 20 samples excluded**
N= 14 insufficient amplifaction in 2D culture
N= 3 contaminations
N=3 lack of adhesion to wells
**52 lung samples**
used for data analysis

**Figure S1**: Flow chart

2D: 2 dimension, g: gram

A

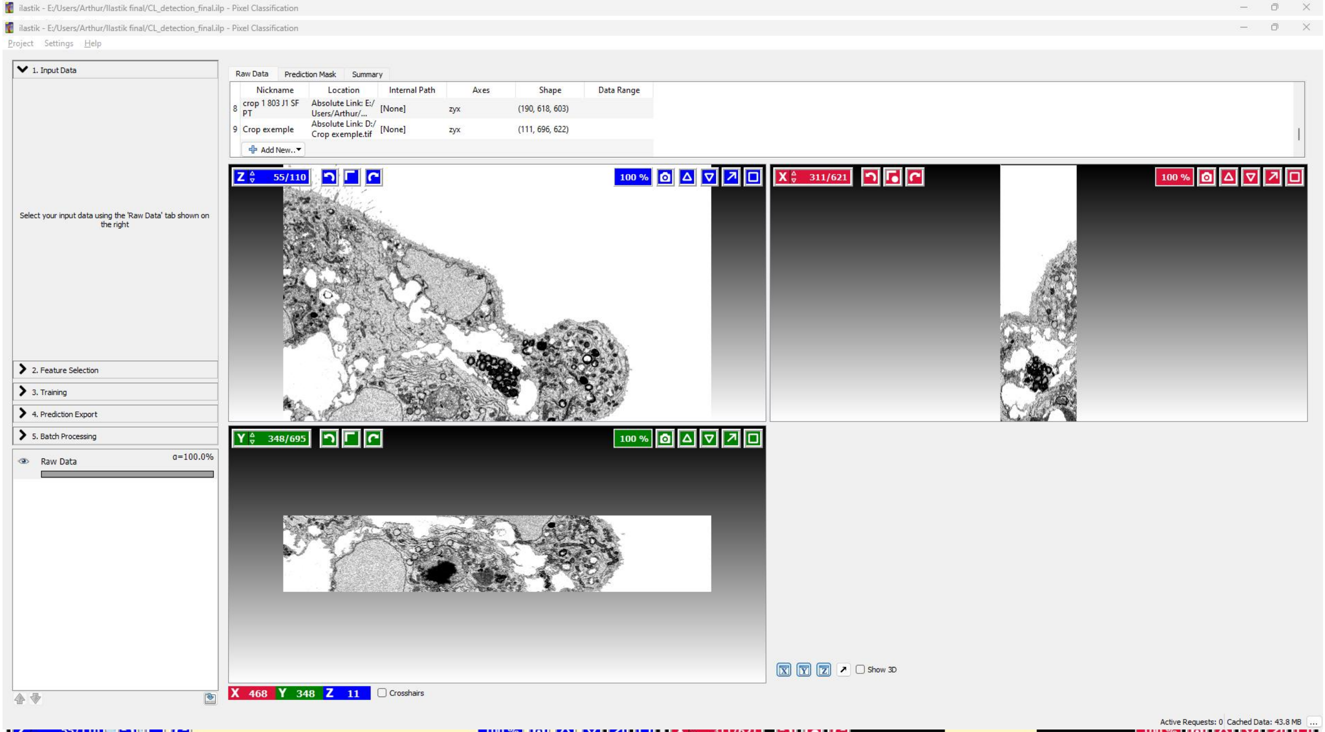
ilastik - E:/Users/Arthur/Ilastik final/CL_detection_final.ilp - Pixel Classification
Project Settings Help
1. Input Data
Raw Data Prediction Mask Summary
Nickname Location Internal Path Axes Shape Data Range
8 crop 1 803 J1 SF PT Absolute Link: E:/ Users/Arthur/... [None] zyx (190, 618, 603)
9 Crop exemple Absolute Link: D:/ Crop exemple.tif [None] zyx (111, 696, 622)
Add New...
Select your input data using the 'Raw Data' tab shown on the right
2. Feature Selection
3. Training
4. Prediction Export
5. Batch Processing
Raw Data α=100.0%
Z 55/110
100 %
X 311/621
Y 348/695
X 468 Y 348 Z 11
Crosshairs
Show 3D
Active Requests: 0 Cached Data: 43.8 MB

B

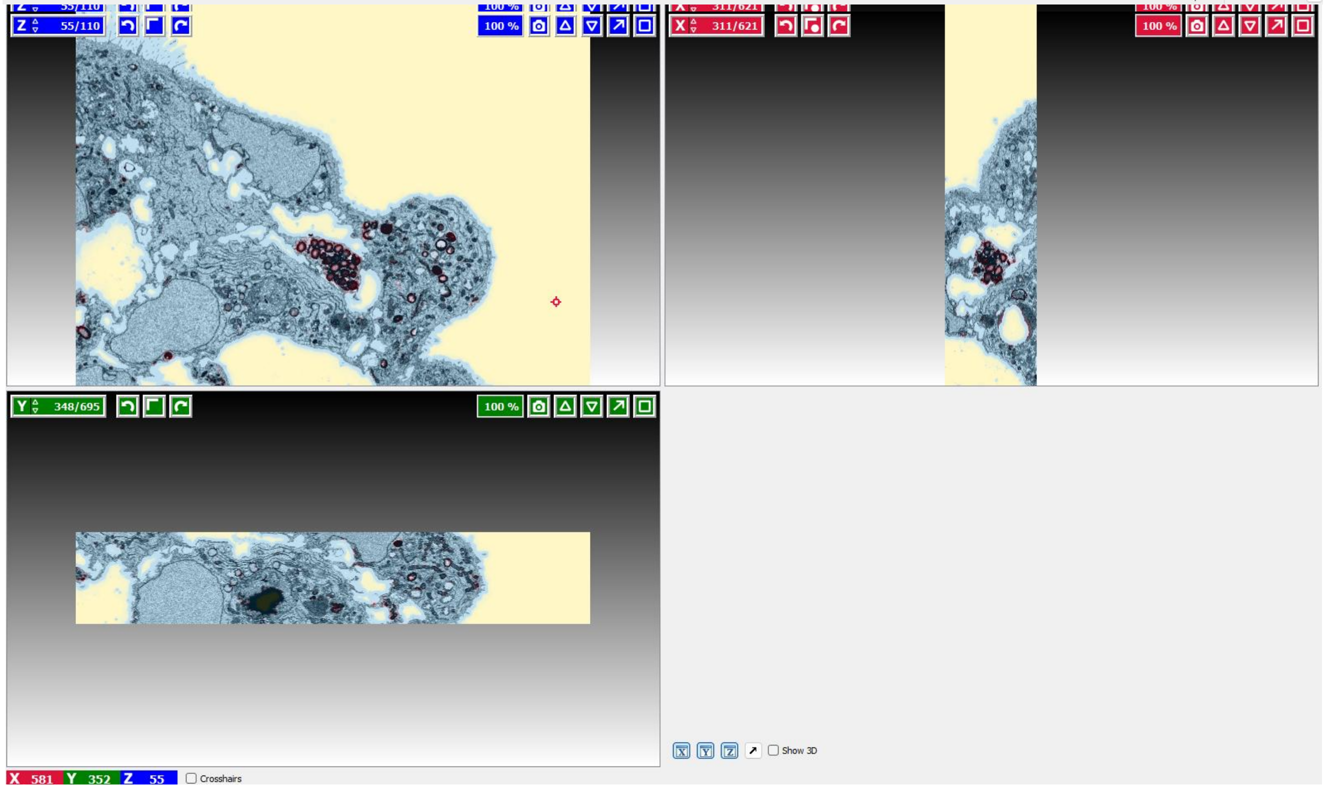
Z 55/110
100 %
X 311/621
Y 348/695
X 581 Y 352 Z 55
Crosshairs
Show 3D

C

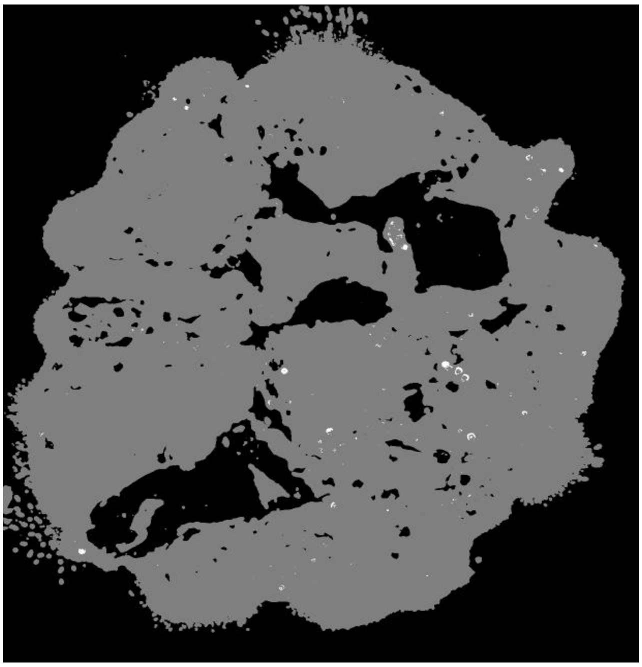

D

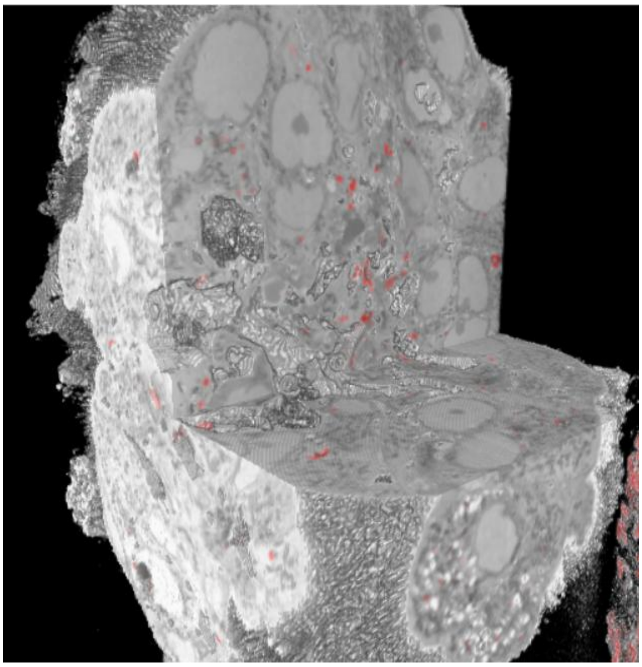

**Figure S2**: **Lamellar bodies (LB) quantification by artificial intelligence (AI) with transmission electron microscopy (TEM) serial block face (SBF) images**
The processing of electron microscopy images to classify organoid cells based on their lamellar body (LB) content was performed following a three-step protocol: training the Ilastik model, training the Cellpose model, and analyzing lamellar body density.

A/ Crops in three, tagging the following structures: "Background," "Cell," and "Lamellar Bodies"
B/ Recognition of lamellar bodies automatically after training cell pose©.
C/ The final mask obtained with the complete stack.
D/2-D reconstruction,
E/ 3-D reconstruction of serial bloc face scanning electron microscopy (SBF-SEM) with artificial intelligence lamellar bodies recognition

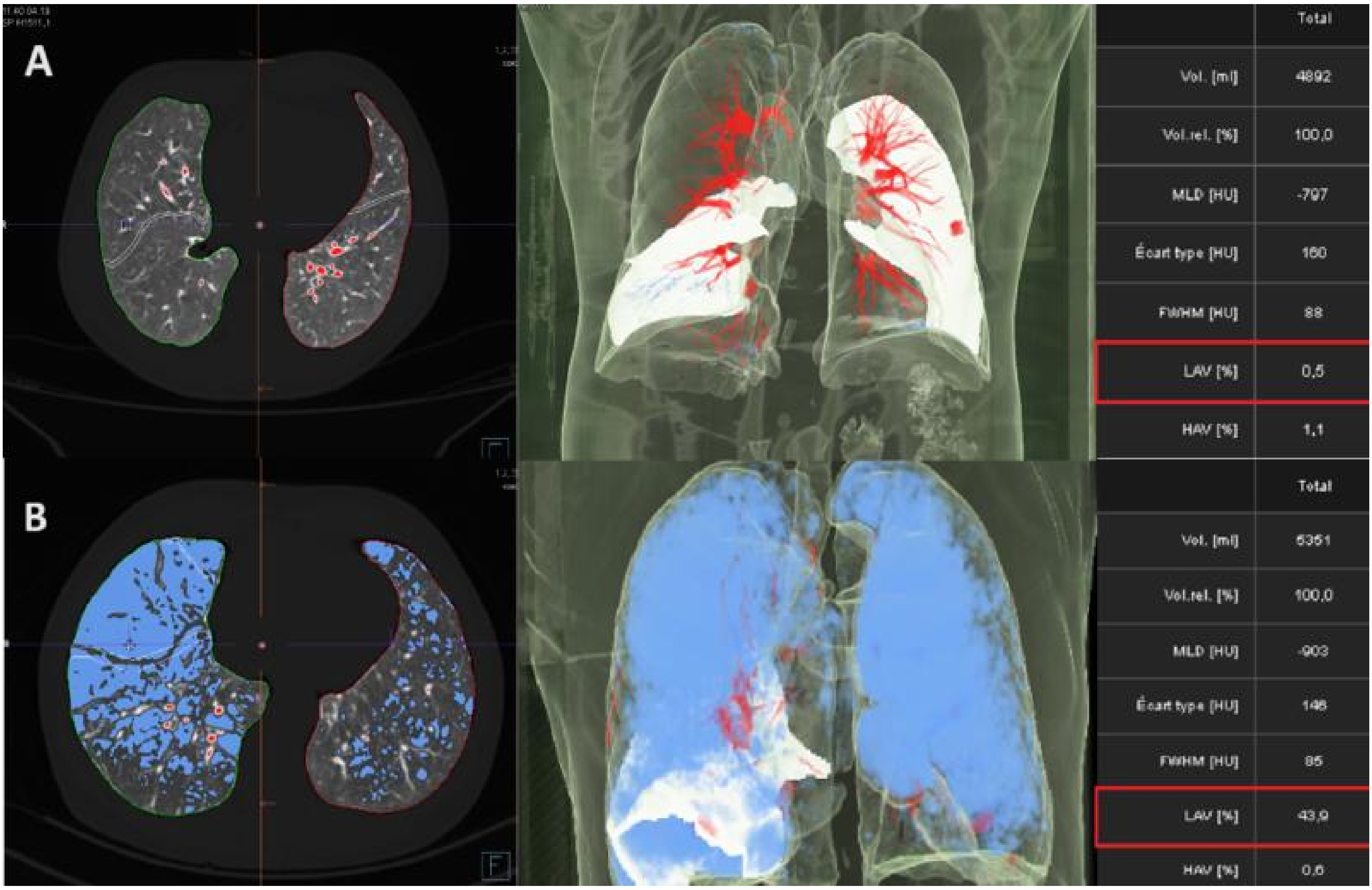
A
Total
Vol. [ml]
4892
Vol.rel. [%]
100,0
MLD [HU]
-797
Écart type [HU]
160
FWHM [HU]
88
LAV [%]
0,5
HAV [%]
1,1
B
Total
Vol. [ml]
6351
Vol.rel. [%]
100,0
MLD [HU]
-903
Écart type [HU]
146
FWHM [HU]
85
LAV [%]
43,9
HAV [%]
0,6

**Figure S3**: Illustration of 3D lung reconstruction and emphysema quantification on chest computed tomography scan using Syngovia© software. A) A non-emphysematous patient (LAA 0.5%) and B) A patient with severe pulmonary emphysema (LAV 43.9%).

FWHM: Full Width at Half Maximum, HAV: Hounsfield Attenuation Value, LAV: Low Attenuation Volume percentage, MLD: Mean Lung Density

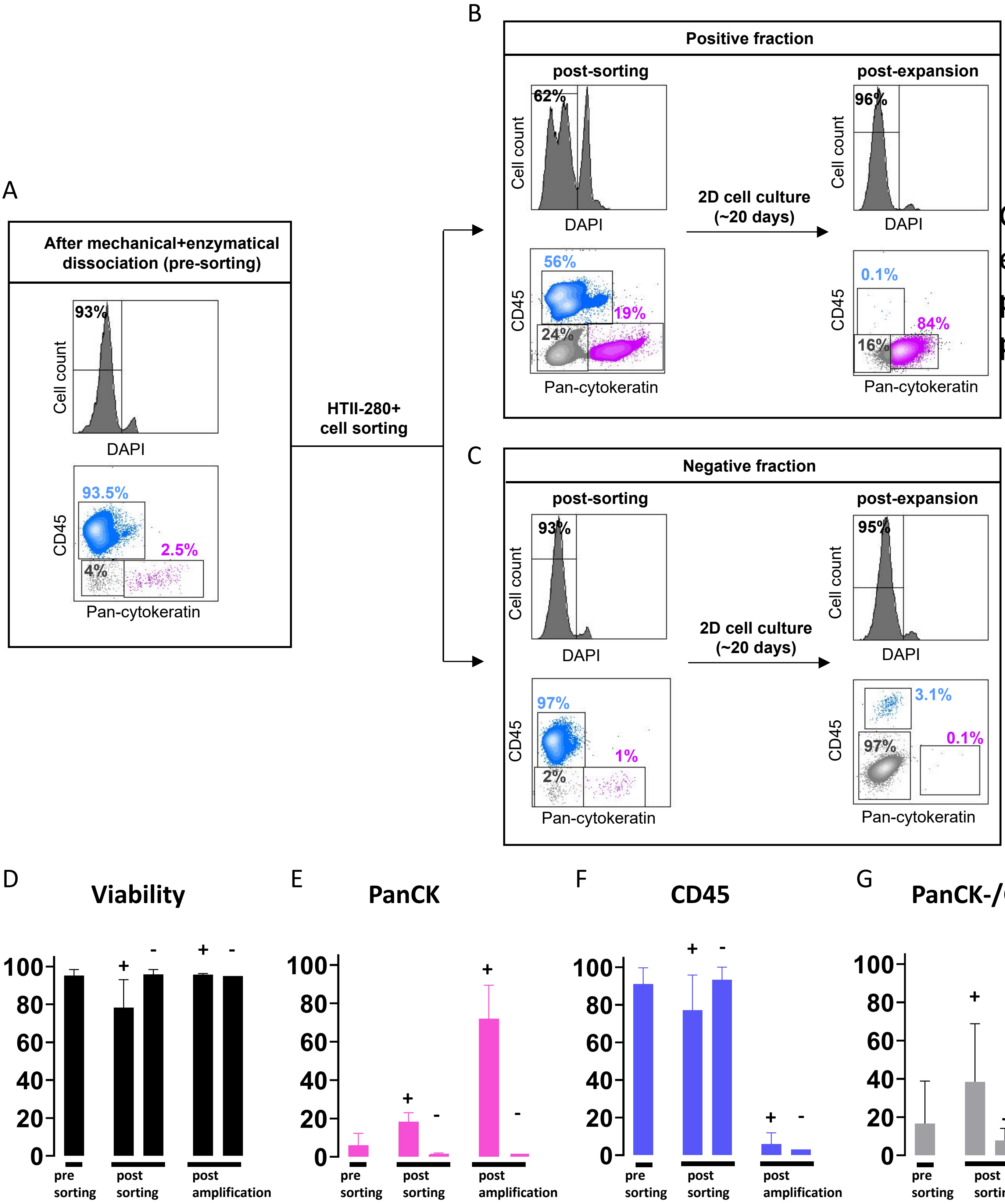


Gating en englobant plus de panck

**Figure S4**: **2D culture characterization by flow cytometry, before microwells seeding**

(A) Flow cytometry after mechanical and enzymatical dissociation (pre-sorting) showing 93% viable cells, 93.5% hematopoietic cells and 2.5% epithelial cells
(B) Flow cytometry after HTII-280 + cell sorting showing 62% viable cells, 56% hematopoietic cells and 19% epithelial cells increasing to 84% after 20 days expansion in 2D culture condition
(C) Flow cytometry after HTII-280 - cell sorting showing 93% viable cells, 97% hematopoietic cells and 1% epithelial cells
(D) Viability of cells after mechanical and enzymatical dissociation, after HTII-280 cell sorting and after 20 days expansion in 2D culture condition (n=3)
(E) Proportion of epithelial cells (panCK+) after mechanical and enzymatical dissociation, after HTII-280 cell sorting and after 20 days expansion in 2D culture condition (n=3)
(F) Proportion of hematopoietic cells (CD45+) after mechanical and enzymatical dissociation, after HTII-280 cell sorting and after 20 days in 2D culture condition (n=3)
(G) Proportion of non epithelial non mononuclear cells (panCK-CD45-) after mechanical and enzymatical dissociation, after HTII-280 cell sorting and after 20 days expansion in 2D culture condition (n=3)

2D: 2 dimension, CD: cluster differentiation, DAPI: 4',6-diamidino-2-phenylindole, HTII-280: Human type 2 cells- 280kDa protein, panCK: pancytokeratin

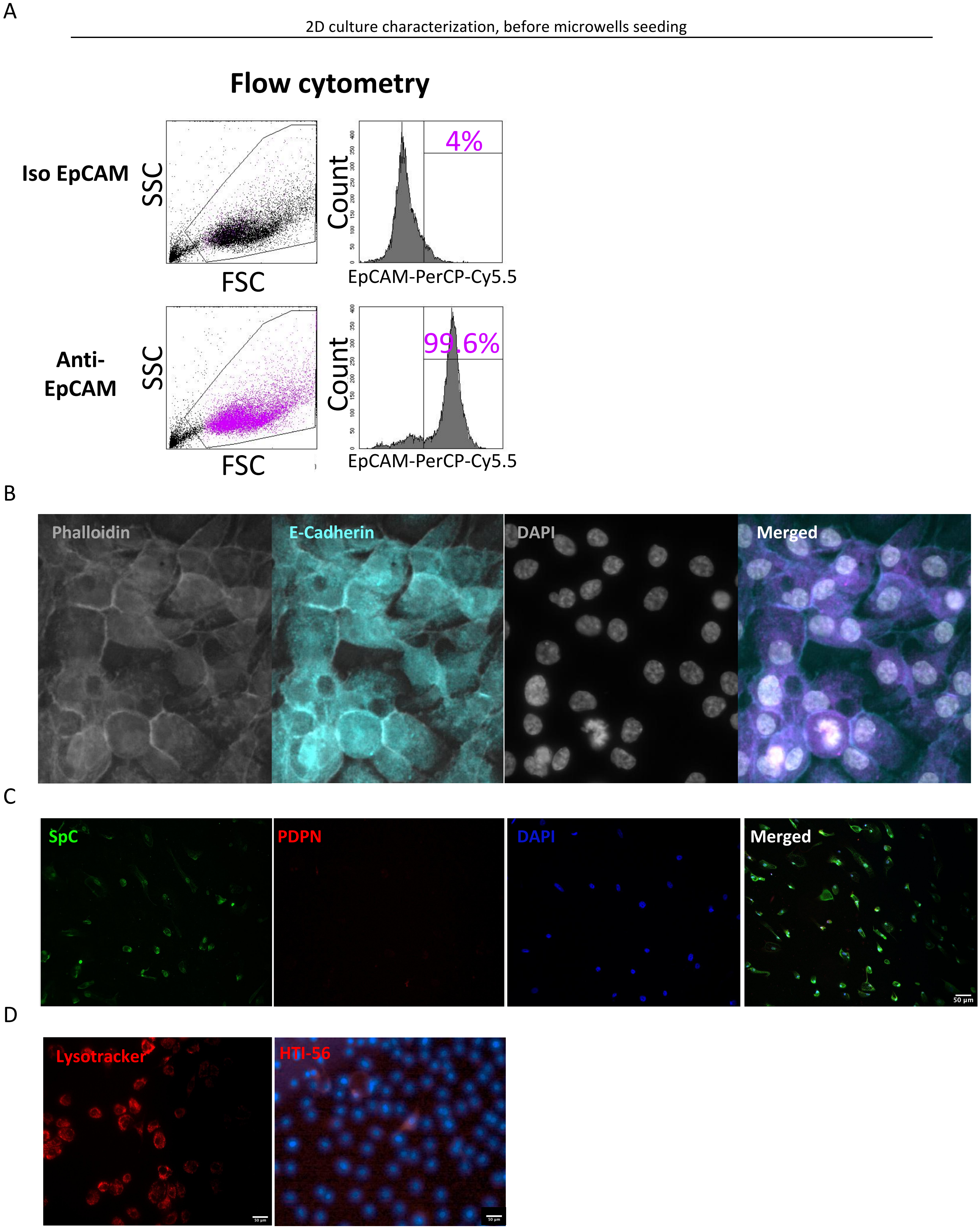
A
2D culture characterization, before microwells seeding
Flow cytometry
Iso EpCAM
SSC
FSC
Count
4%
EpCAM-PerCP-Cy5.5
Anti-EpCAM
SSC
FSC
Count
99.6%
EpCAM-PerCP-Cy5.5
B
Phalloidin
E-Cadherin
DAPI
Merged
C
SpC
PDPN
DAPI
Merged
50 µm
D
Lysotracker
HTI-56

**Figure S5**: **2D culture characterization, before microwells seeding (~ 20+/-5 days after 2D seeding)**

(A) fluorescent image of 2D-culture stained for cystoskeleton (phalloidin, grey), epithelial junction (E-cadherin, blue) and nuclei (DAPI, white) confirming epithelial phenotype
(B) fluorescent image of 2D-culture stained for AEC2 (Spc, green), AEC1 (podoplanin, red) and nuclei (DAPI, blue) confirming a majority of AEC2
(C) fluorescent image of 2D-culture stained for AEC2 (lysotracker, red), AEC1 (HTI-56, red) confirming a majority of AEC2
(D) Flow cytometry showing 96% of epithelial phenotype (epCAM +) 20 days after 2D seeding

2-D: 2 dimension, AEC1: alveolar type 1 cell, AEC2: alveolar type 2 cell, DAPI: 4',6-diamidino-2-phenylindole, EpCAM: Epithelial cell adhesion molecule, , HTI-56: Human type 1 cells-56kDa protein, PDPN: podoplanin, SPC: surfactant protein C

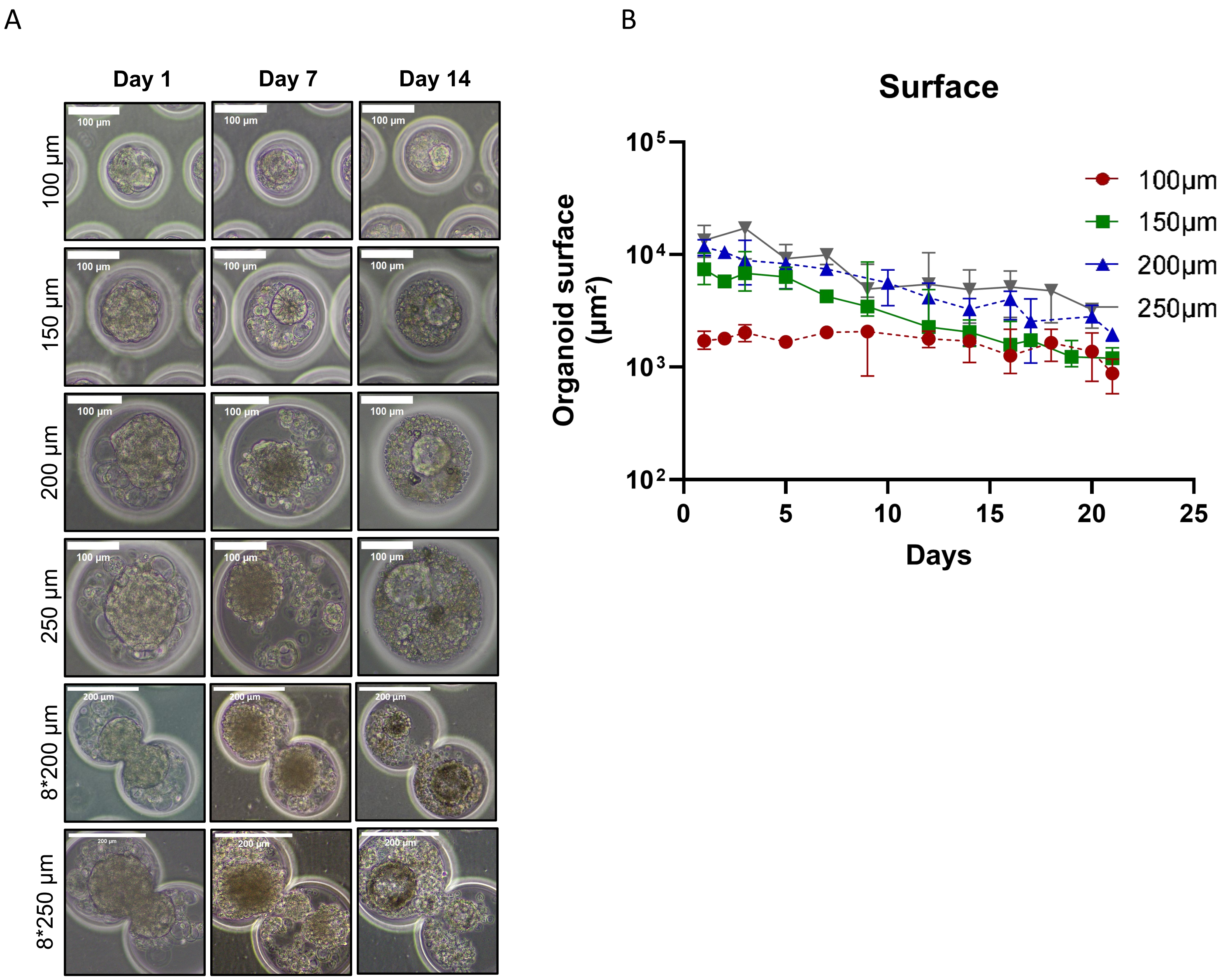

A
B
Day 1
Day 7
Day 14
100 µm
150 µm
200 µm
250 µm
8*200 µm
8*250 µm
Surface
Organoid surface (µm²)
Days
100µm
150µm
200µm
250µm

**Figure S6**: 3D alveolosphere model from human type 2 alveolar epithelial cells.

(A) Alveolosphere maintained in culture until D14 in different diameters (100, 150, 200 and 250µm) shapes (circle and double circle) of microwells.

(B) Alveolospheres surface until D14 according to different diameters (100, 150, 200 and 250µm). Dotted lines relate to data already shown in Figure 1G

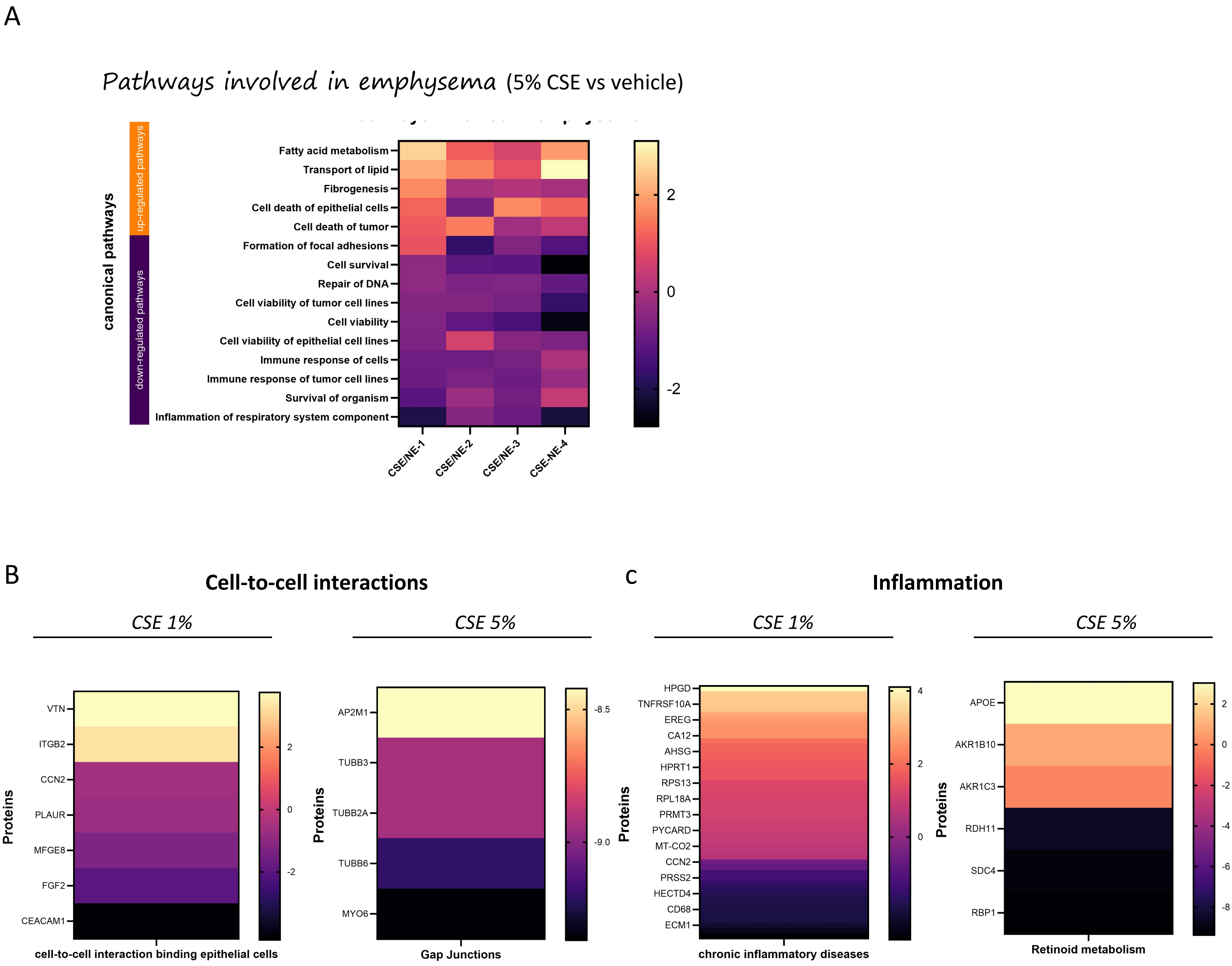
A
Pathways involved in emphysema (5% CSE vs vehicle)
canonical pathways
up-regulated pathways
down-regulated pathways
Fatty acid metabolism
Transport of lipid
Fibrogenesis
Cell death of epithelial cells
Cell death of tumor
Formation of focal adhesions
Cell survival
Repair of DNA
Cell viability of tumor cell lines
Cell viability
Cell viability of epithelial cell lines
Immune response of cells
Immune response of tumor cell lines
Survival of organism
Inflammation of respiratory system component
CSE/NE-1
CSE/NE-2
CSE/NE-3
CSE-NE-4
2
0
-2
B
Cell-to-cell interactions
CSE 1%
CSE 5%
Proteins
VTN
ITGB2
CCN2
PLAUR
MFGE8
FGF2
CEACAM1
2
0
-2
cell-to-cell interaction binding epithelial cells
AP2M1
TUBB3
TUBB2A
TUBB6
MYO6
-8.5
-9.0
Gap Junctions
C
Inflammation
CSE 1%
CSE 5%
HPGD
TNFRSF10A
EREG
CA12
AHSG
HPRT1
RPS13
RPL18A
PRMT3
PYCARD
MT-CO2
CCN2
PRSS2
HECTD4
CD68
ECM1
4
2
0
chronic inflammatory diseases
APOE
AKR1B10
AKR1C3
RDH11
SDC4
RBP1
2
0
-2
-4
-6
-8
Retinoid metabolism

**Figure S7**: ***Emphysema modelling in the 3D-alveolosphere model by exposure to 5% cigarette smoke extract (CSE)***

(A) Heatmap of the differentially expressed proteins between 5% CSE and vehicle exposed alveolospheres. Five up-regulated and ten down-regulated major pathways involved in emphysema were identified, based on IPA (orange: activation; purple: inhibition).

(B) Heatmap of six differentially expressed proteins between CSE and vehicle exposed alveolospheres; most of them being down-regulated (epithelial cellular junction, remodeling epithelial adherens, pulmonary signaling healing pathway, cytoskeleton signaling pathway, cellular response to hypoxia) and some being up-regulated (airway inflammation) (orange: activation; purple: inhibition).

(C) Heatmap of six differentially expressed proteins between CSE and vehicle exposed alveolospheres; most of them being down-regulated (epithelial cellular junction, remodeling epithelial adherens, pulmonary signaling healing pathway, cytoskeleton signaling pathway, cellular response to hypoxia) and some being up-regulated (airway inflammation) (orange: activation; purple: inhibition).

3-D: 3 dimension, *CSE: cigarette smoke extract*
